# Spatial preferences account for inter-animal variability during the continual learning of a dynamic cognitive task

**DOI:** 10.1101/808006

**Authors:** David B. Kastner, Eric A. Miller, Zhounan Yang, Demetris K. Roumis, Daniel F. Liu, Loren M. Frank, Peter Dayan

## Abstract

Individual animals perform tasks in different ways, yet the nature and origin of that variability is poorly understood. In the context of spatial memory tasks, variability is often interpreted as resulting from differences in memory ability, but the validity of this interpretation is seldom tested since we lack a systematic approach for identifying and understanding factors that make one animal’s behavior different than another. Here we identify such factors in the context of spatial alternation in rats, a task often described as relying solely on memory of past choices. We combine hypothesis-driven behavioral design and reinforcement learning modeling to identify spatial preferences that, when combined with memory, support learning of a spatial alternation task. Identifying these preferences allows us to capture differences among animals, including differences in overall learning ability. Our results show that to understand the complexity of behavior requires quantitative accounts of the preferences of each animal.

## Introduction

Modeling animal behavior provides a rigorous and falsifiable way to formulate quantitatively the computations that underlie the decisions that drive actions. Such modeling is often applied to tasks under tight experimental control via momentary sensory input or motor output, and the models are typically created to capture the behavior after acquisition of the task has occurred. These detailed models can capture much of the variability of animal behavior, and have thereby provided variables that help explain neural activity patterns recorded during these tasks^1–5^.

Many tasks are not under such tight experimental control, however, and often the goal is to understand not asymptotic performance but instead the learning process whereby experience drives systematic changes in behavior. Such learning and memory tasks, which include the Morris water maze^6–8^, the Barnes maze^9–11^, the T-maze^12, 13^ and the W-track^14–17^, are widely used, but rigorous and systematic models of these behaviors are less common as methods of understanding the actions of the animals. Instead, behavior in these tasks is interpreted using intuition and qualitative, model-agnostic, metrics that are intended to capture the underlying learning and memory processes.

We have recently shown that such intuition is not sufficient to capture the course of learning in a simple spatial alternation task: a simple agent endowed with perfect spatial memory could not learn the task as quickly as an animal^18^. This result calls into question the common process of interpreting spatial alternation behavior only in terms of memory. In that work, we hypothesized that pre-existing or trainable preferences for particular locations or transitions between locations, in addition to memory, could underlie the learning of spatial tasks; however, the structure of the simple task limited any validation of that hypothesis. Here we sought to develop a systematic approach to determine the computational components for learning spatial alternation. We sought a solution that would provide critical information about the variables that best describe the different behavior of individuals, as well as the whole group. We expect that this would allow accurate inferences about individually idiosyncratic neural activity patterns along with the algorithms implemented by those patterns.

Modeling the entire course of learning is central to this approach; however, modeling the entire course of learning is difficult for experimental and analytical reasons. Experimentally, animals are typically exposed to behavioral tasks without prior experiences that would aid in identifying, or controlling, pre-existing preferences. In particular, these tasks are often implemented by combining the learning about the spatial environment in which the task is set with learning of the rules of the task. This makes it challenging to quantify pre-existing preferences that manifest in the way in which an animal interacts with the space independent of the constraints of the task. Moreover, at least until recently^19–22^, it has been common to shape the behavior of each subject in an individualized and heterogeneous manner. This is well suited to studies of asymptotic performance of tasks, but problematic for understanding the acquisition of those tasks.

Analytically, each subject only provides one set of data points about their entire learning trajectory. This makes it challenging to convincingly fit a model to individual animal behavior, and attribute that single learning course to innate capabilities as opposed to random variation. The standard implementations of Reinforcement Learning (RL)^23^ models that capture these behaviors also presents challenges. RL formalizes the notion of updating information about a task based upon actions taken and rewards delivered and has the capacity to learn highly complex tasks^24–27^. However, the standard RL agents that capture asymptotic adaptivity extremely well often fail to describe competently the whole course of learning. In general, agents learn more slowly than animals do, since the agents do not embody the extensive knowledge about the world that animals apparently possess. Specific, model-based agents, by contrast, can be constructed to absorb the relevant information efficiently (or be provided with an advantageously restricted set of inputs). These models (unfairly) gain information from the outset as to which are the critical features of the environment^28, 29^ and thus learn very quickly. In sum, neither standard RL nor model-based RL, as normally implemented, matches the learning rates of the animals.

Here we present a general approach that addresses these challenges. We develop an automated behavioral system that minimizes tailored shaping and provides a higher-throughput method to record behavior. This allows for more animals and more data for model validation. We include a period of free exploration prior to beginning an alternation task, allowing for the measurement of unconstrained behavioral preferences of each individual animal. We design an extended alternation task that includes a series of increasingly complex alternation contingencies providing multiple learning opportunities for each animal, thereby allowing for a distinction between random variation and innate capacity. Finally, we develop a series of RL agents specifically focused on capturing the dynamics of learning of individual animals. The result is a quantitative understanding of spatial alternation behavior that concludes that not only is memory critical for the way in which rats perform a spatial alternation task, but also that dynamic preferences play a large role in determining the choices that individual animals make.

## Results

### Automated system for rats to learn a series of spatial alternation contingencies

As our goal was to model the entire course of learning, and thereby understand the computations that underlie spatial alternation behavior, it was critical to standardize the behavioral training and to reduce potential effects of experimenter-subject interactions on learning^30^. We therefore developed an automated behavioral system^19–21^ that requires minimal animal handling: once animals were placed in the apparatus, no further experimenter contact was necessary until the end of the daily behavior. This system also enables the measurement of behavior across many animals throughout the entire course of learning and performance of the task. The apparatus contains four parts: 1) a six-armed track with reward wells at the end of each arm; 2) four rest boxes, each with a reward well; 3) corridors connecting the rest boxes to the track; and 4) doors to gate the pathway on and off the track for each rest box (Fig. 1A).

**Figure 1.**
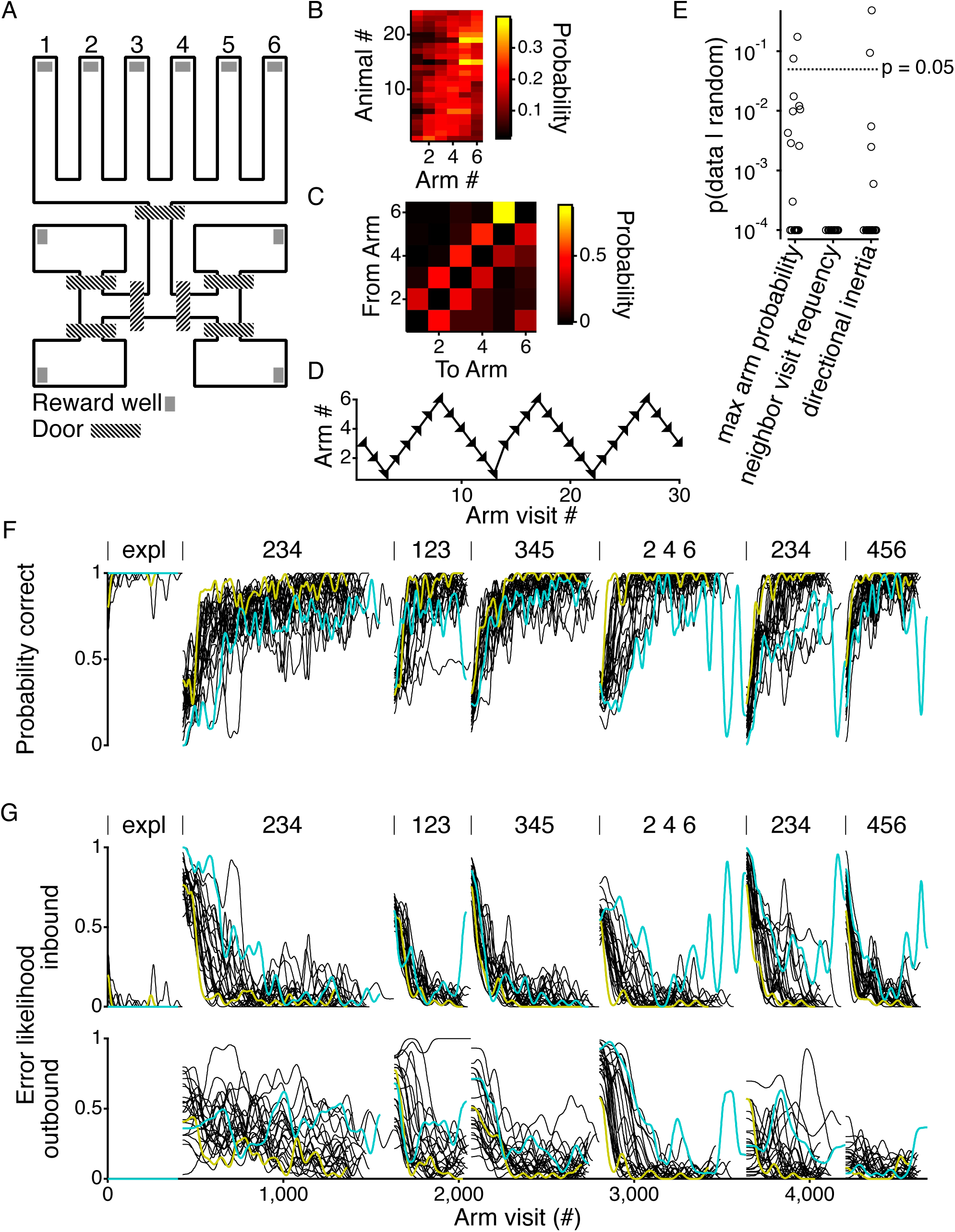
Automated behavior system for analysis of continuous spatial alternation behavior. (A). Layout of automated behavior system. (B) Arm preferences of all rats (*n* = 24) during the exploratory period of the behavior, where a rat can get rewarded at any arm of the track. Rats ordered by arm preference. (C) Example transition matrix during the exploratory period of the behavior a single rat (animal #8 from A) showing the probability of going to any of the six arms when starting from each of the six arms. (D) Example arm choices (arrowheads) of a single rat (animal #1 from A) during a session of the exploratory behavior. (E) Probability of seeing the maximal arm preference (left) neighbor visit frequency (middle) or directional inertia (right) given random choices between the six arms. Horizontal line shows a probability of 0.05. For the neighbor transition frequency, the random distribution was defined by the arm visit probability of the animal, and for the directional inertia the random distribution was defined by the transition matrix of the animal. As the p-value was determined using 10,000 draws from distributions, the minimal value is 10^−4^. 14/24 rats were at that minimal value for the max arm probability, 24/24 for the neighbor visit frequency, and 19/24 for the directional inertia. (F) Probability of getting a reward for all 24 rats. Within each contingency, curves smoothed with a Gaussian filter with a standard deviation of 10 arm visits. Two different rats shown in colors (yellow and teal) to indicate consistency of performance in those rats across the different contingencies. Beginning of each contingency is demarcated by vertical lines above the plot. Contingencies indicated by the 3 arms that have the potential to be rewarded. (G) Error likelihoods for inbound and outbound trials for all 24 animals. Values smoothed with a Gaussian filter with a standard deviation of 10 inbound or outbound trials and then interpolated to reflect total arm visits. Colors indicate the same rats as in F. Contingencies indicated as in F.

Our previous work suggested that accurate descriptions of learning might require dynamic preferences, defined as (changeable) tendencies for animals to prefer specific locations or specific transitions between locations^18^. It was therefore critical to measure the initial values for these preferences. Furthermore, we sought to disambiguate the learning of the task from the learning of the space of the task. Therefore prior to the rats beginning the spatial alternations task, they had 14 -16 sessions (362 -425 total trials) of exploration on the track wherein the rats were rewarded at any arm visited as long as it was not a repeat visit to the immediately preceding arm.

These exploration sessions revealed multiple preferences. First, individual rats showed preferences towards visiting specific arms (Fig. 1B&E). 22 of the 24 rats showed significant (*p* < 0.05) deviation from a random arm visit pattern (*p* = 1.1 × 10^−6^, see Methods). Second, rats also had a large propensity to transition from their current arm to neighboring arms (Fig. 1C&E), with all 24 rats showing significant (*p* < 0.05) deviation from randomly transitioning between arms, even given their individual arm visit probabilities (*p* = 6.3 × 10^−8^, see Methods). And finally, the rats exhibited directional inertia, calculated as the frequency of an animal going in the same direction as it did on the immediately preceding trial (Fig. 1D&E). A partially different 22 out of 24 rats showed significant (*p* < 0.05) deviation from random directional inertia, even accounting for their individual transition probabilities (*p* = 1.7 x 10^−6^, see Methods).

Given the existence of these preferences, we proceeded to ask whether those preferences play a role in learning. Following this initial exploratory period, and without any external signal to indicate a change, rats were sequentially exposed to different spatial alternation contingencies. The six arms of the track allow for the learning of multiple spatial alternation contingencies^31, 32^, and we exposed animals to six different contingencies to provide multiple learning exposures with different levels of difficulty (Fig. 1F). These exposures in turn help constrain the models for each individual animal and provide opportunities to cross-validate the models by training on one set of contingencies and testing on another.

In each contingency, only three arms had the potential to deliver reward. Reward was delivered within a given contingency if the rat alternates between the outer arms after every visit to the center arm. For instance, if the contingency was at arms 2-3-4, then to get reward the animal would have to follow the sequence 3-4-3-2-3-4-3 etc. Following previous studies in a related environment^15, 33^ we defined inbound trials as trials where the animal starts from an arm that is not the center arm (arm 3 in this example), and outbound trials as trials where the animal starts at the center arm of the contingency.

Performance improved on each of the contingencies, such that by the end of each one, rats typically made few outbound or inbound errors (Fig. 1G, S1B). There was, however, substantial and systematic variability across animals, where individual animals consistently showed higher or lower performance across all contingencies (see yellow and cyan colored lines in Fig. 1F&G for examples). This variability provided an additional goal for our modeling, in that an ideal model would capture not only the overall learning of the group but also the differences among individuals.

### Modeling framework

In this work we use a similar modeling framework as our previous study^18^. For clarity and completeness, we describe and motivate the choices of that model. We first specify the algorithm that we use at the base of the model. We use a simple algorithm that, like the animals, does not require acausal information, can alter its internal information based upon its choices and rewards to increase the expected return of reward, and can work in the face of partial observability (see below). This led us to the actor-critic class of RL accounts trained by the REINFORCE policy gradient algorithm^34^, and employing a form of working memory^35^. REINFORCE is a popular choice for characterizing animal learning behavior in RL paradigms^36^, and there is also evidence of its use in humans^37^.

Given that algorithm, we can specify a family of models with a common form. The models describe the behavior of an agent choosing an arm on trial *t*, which we write as at. The choice of at depends probabilistically on an internal characterization of its situation or state *s_t_*, which can contain various sorts of information such as past arm choices. This dependence arises through a collection of action preferences or propensities *m*(*a*, *s_t_*), such that actions with higher propensities are more likely to be chosen. The propensities are updated as a function of reward. The full details of the equations involved are provided in the Methods. In brief: a conventional softmax function converts the propensities to probabilities, *p*(*a*; *s_t_*), of choosing to go to arm *a_t_*_+1_ = *a* on this trial (Eq. 1). Via the rules of the task, this choice of arm then determines whether the model receives a reward, *r_t_*_+1_, and also causes the state to update to *s_t_*_+1_. This reward is then used to calculate the prediction error, *δ*t, using the value function of the critic at states *s_t_* and *s_t_*_+1_, *V*(*s_t_*) and *V*(*s_t_*_+1_) (Eq. 2). *δ_t_* is then used to update *V*(*s_t_*) (Eq. 6) and the factors governing the propensities *m*(*a*, *s_t_*) (Eq. 3 – 5). Finally, new propensities *m*(*a*,*s_t_*_+1_) are calculated, at which point the process begins again with the agent choosing its next arm to visit (Fig. 2A&B).

**Figure 2.**
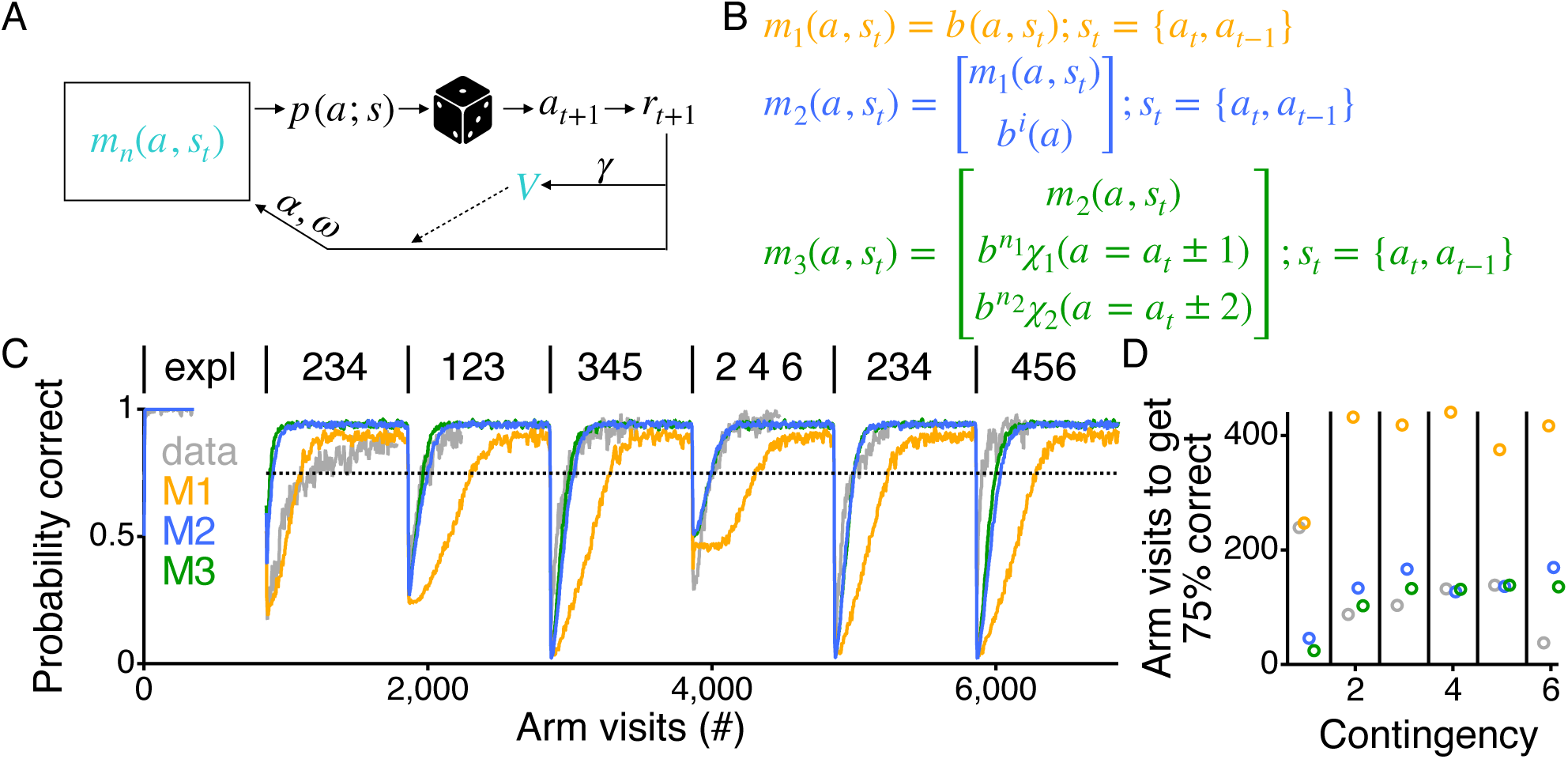
RL model with working memory and dynamic preferences can learn as rapidly as the rats. (A) Graphic of RL agent. Colored symbols, *m*_*n*_(*a*,*s*_*t*_) and *V*, indicate the components that change as the agent goes to arms, *a*, and does or does not get reward, *r*. (B) The different components of the propensities, *m*_*n*_(*a*,*s*_*t*_), for the different models. The state of the agent, and therefore the probability of transitioning to each of the arms, *p*(*a*; *s*), is defined by either just the current arm location, *a_t_*, or by the current arm location, *a_t_*, and the previous arm location, *a_t_*_−1_, of the agent. *b^i^*(*a*) is the independent arm preference. 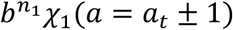 and 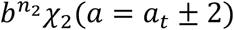 are the preference to transition to a neighbor 1 or 2 arms away, respectively. (C) Average reward probability of all animals (n = 24) across all contingencies (grey), and average behavior of 200 repeats of the models with parameters chosen to maximize the rewards received across all contingencies. The model was given extra arm visits to reach asymptotic behavior (after the endpoints of the grey curves for each contingency) to show more clearly the model’s ability to learn the task. Dotted horizontal lines show 75% probability correct. Contingencies indicated as in Fig 1F. (D) Number of trials to pass 75% probability correct for the data (grey) and models; however, M1 not shown as it never performs above 75% correct.

All of the models described below have only three parameters, all of which take values between 0 and 1. The first parameter is the temporal discount factor, *γ*, which determines the weighting of rewards in the farther future in defining the long-run values of states (and thus in calculating the prediction error, *δ*) (Eq. 2). The second parameter is the learning rate, *α*, which determines how much *δ* updates the propensities and the value function (Eq. 3–6). The third parameter is the forgetting rate, *ω*, which determines how quickly the propensity parameters and the value function decay towards 0 (Eq. 3–6), a value that would indicate that there is no specific information about which arm to visit in any state since the value would be the same for all propensities. *ω* enables the model to adapt to the nonstationarity of the task by constantly depreciating old information, allowing for changes in the propensities and the value function during changes in contingencies.

The framework described above falls into the category of model free (MF) RL agents and MFRL agents typically learn slower than animals. Therefore, to develop a model that has the potential to learn as quickly as individual rats continually learn a task, we started by comparing the best each model could do to the average behavior across all rats (Fig. 2C&D). This provided a straightforward way to determine if the model had the potential to fit individual animals because if the best version of the model could not learn as quickly as the animals there would be no chance for it to capture the learning of all of the individual animals.

Finally, we note that our goal was not to perfectly recapitulate all aspects of each animal’s behavior, as such a task is well beyond our current understanding. Instead, we sought to develop a simple, interpretable model that could capture learning rates across at least a subset of contingencies. Such a model would allow us to determine whether incorporating spatial preferences was important for describing behavior. That model, if it could be fit to individual animals, could also help us quantify differences in behavior among individuals.

Finally, areas of lack of fit would provide a clear direction forward for future augmentation and understanding.

### Memory alone is not sufficient

We previously demonstrated that a model with working memory alone does not capture the rapid way rats learn a simple spatial alternation task^18^. We replicate and extend that finding in this more complex environment using our first model (M1). As we did in the previous work^18^, we added a memory component following an approach by Todd et al.^35^ where the state of the model is augmented to include a memory unit that stores the immediate past action. This enables the models to make decisions based upon current and past information. Such a strategy has been used to learn common rat behavioral tasks^38^, and exhibits features of rat behavior^39^. In all of the models, the state, *s*_*t*_ = {*a*_*t*_, *a*_*t*−1_}, includes both the current and the most recent past arm (Fig. 2B). For model M1 the propensities are *m*_1_(*a*,*s*_*t*_) = *b*(*a*|*a*_*t*_, *a*_*t*−1_. For each state, *b*(*a*|*a*_*t*_, *a*_*t*−1_ contains 5 numbers governing the propensity to make a transition from the current arm to each of the other 5 arms. Returning to the same arm is not allowed in the model, as it was never rewarded in the behavior.

This working memory (WM) RL agent starts with perfect memory of the immediate past and has the capacity to perform each contingency well; however, it learns to do so far slower than the average across the rats (Fig. 2C&D), even when the parameters are set to optimize the obtained reward. With M1, good performance on the first contingency arises at the correct timescale—something that will be discussed further below—but performance on all the subsequent contingencies improves much more slowly than for the rats. For contingencies 2 -5, M1 reached 75% correct 2.7 -4.9 times slower than the average performance of the rats, and for contingency 6 M1 was 10.9 times slower.

### Arm and transition preferences, combined with memory, enables the model to learn as rapidly as the rats

Given the failure of M1 to show relevant learning rates, we then asked whether the incorporation of dynamic preferences would be sufficient to enable rapid learning, as was the case for the simpler three-arm version of the task^18^. To capture the preferences, we added individual propensities to the model. As the goal of modeling is always to develop the simplest model that explains the data, we begin by adding a single term for each arm to capture the individual arm preferences shown by the animals (Fig. 1B, S2A). This yields model M2, where *m*_2_(*a*,*s*_*t*_) = *b*(*a*|*a*_*t*_, *a*_*t*−1_) + *b*^*i*^(*a*) (Fig. 2B). The term *b^i^* (*a*), which we call a dynamic independent arm preference, provides the agent with additional preferences to choose specific arms next, independent of its current or past locations. As with the state-dependent propensity terms, *b^i^* (*a*) are also updated by *δ*_*t*_ through the process of learning. Importantly, adding this term allows us to capture both the fact that the animals may prefer specific arms before beginning the learning of the alternation contingencies and that these preferences can be dynamic and shaped by reward. Importantly, including this term or any other preference related term does not entail adding any additional free parameter to the model.

Including the dynamic independent arm preference yields an agent that can learn much more quickly, but still failed to match the learning rates of the rats (Fig. 2C&D). M2 learned the first contingency faster than the animals, reaching 75% correct 5 times faster than the rats. By contrast, for contingencies 2 -5, M2 reached 75% correct 1.0 -1.6 times slower than the average performance of the rats, and for contingency 6 M2 was 4.4 times slower.

The failure to match learning rates led us to incorporate an additional preference observed in the animals, a dynamic transition preference. This yielded model M3, for which 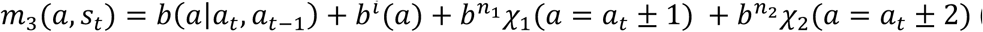 (Fig. 2B, S2B). The additional propensities capture the preference of the animals to transition to neighbors that are either one, 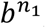, or (as an addition to our previous model,^18^ because of the structure of the task) two arms 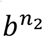, away, independent of the current location of the animal (Fig. 1C). Here again these state values update using the same three parameters as the previous models.

Model M3 more closely approximates the behavior of the rats (Fig. 2C&D). While M3 reached 75% correct 10 times faster than the rats on the first contingency and 3.6 times slower for contingency 6, for contingency 2 -5, M3 reached 75% correct at rates more similar to the average performance of the rats.

### Model with memory and arm and transition preferences fits individual animals

M3, despite its relative simplicity, proved sufficiently flexible to match the average learning rates of the animals for some contingencies. This in turn suggested that it could be sufficiently powerful to capture important aspects of the behavior of individual rats. For the fit to an individual rat, we forced the model to make the same sequence of arm visits as the animal during the initial exploratory phase, effectively using the data of the animal to inform the initial condition of the model. We then fit aspects of selected contingencies, testing how well the resulting parameters permitted generalization to the behavior in the other contingencies.

To determine the best fitting parameters we used an Approximate Bayesian Computation (ABC) method^40^, consistent with other studies using RL agents to fit rodent behavior^39, 41^. ABC methods find parameters such that the average behavior of the model when operating in the task, choosing stochastically, matches as well as possible that of an individual animal, according to some suitably chosen statistics. We averaged 200 repeats of the model and chose as statistics the inbound and outbound performance for the contingencies we fit. We then evaluated the fit of each model to each animal by calculating the root mean square (rms) difference between the model and data on inbound and outbound trials on all contingencies.

We found that even though the model was able to fit to the inbound and outbound errors of the first contingency (Fig. S3A), the parameters from those fits did a poor job of capturing the behavior of the animals on subsequent contingencies (Fig. S3B). This failure was not too surprising given that the first contingency was an outlier when evaluating the maximal reward the models could receive (Fig. 2C). We will return to understand this difference in the first contingency below.

Therefore, we chose to fit specifically the second and third contingencies. These contingencies are the most representative for this task, as 1) both follow other simple contingencies, and 2) occur before the hardest, fourth, contingency, for which the required alternation involves skipping neighboring arms. Fitting the second and third contingencies allowed us to use the performance of the model on the subsequent, and preceding, contingencies as predictions that could test the goodness of fit of the models. Finally, to verify that the additional preferences of M3 were necessary for the fit to individual animals, we also fit to M1 and M2.

The fits of the second and third contingencies (Fig 3A) confirmed that M3 fit the individual animals better than M2 and M1 (Fig. 3B). Specifically, both M2 and M3 fit inbound and outbound errors with lower rms errors as compared to M1 (p < 10^−6^, paired permutation test), and M3 improved upon M2’s performance for outbound errors (*p* = 1.4 × 10^−4^, paired permutation test). These findings indicate that incorporating all three observed propensities— memory, independent arm and neighbor transitions preferences—improves the fit of the model to the data. We note that there remain clear situations in which M3 still does not fit the data well, however. We return to this observation below.

**Figure 3.**
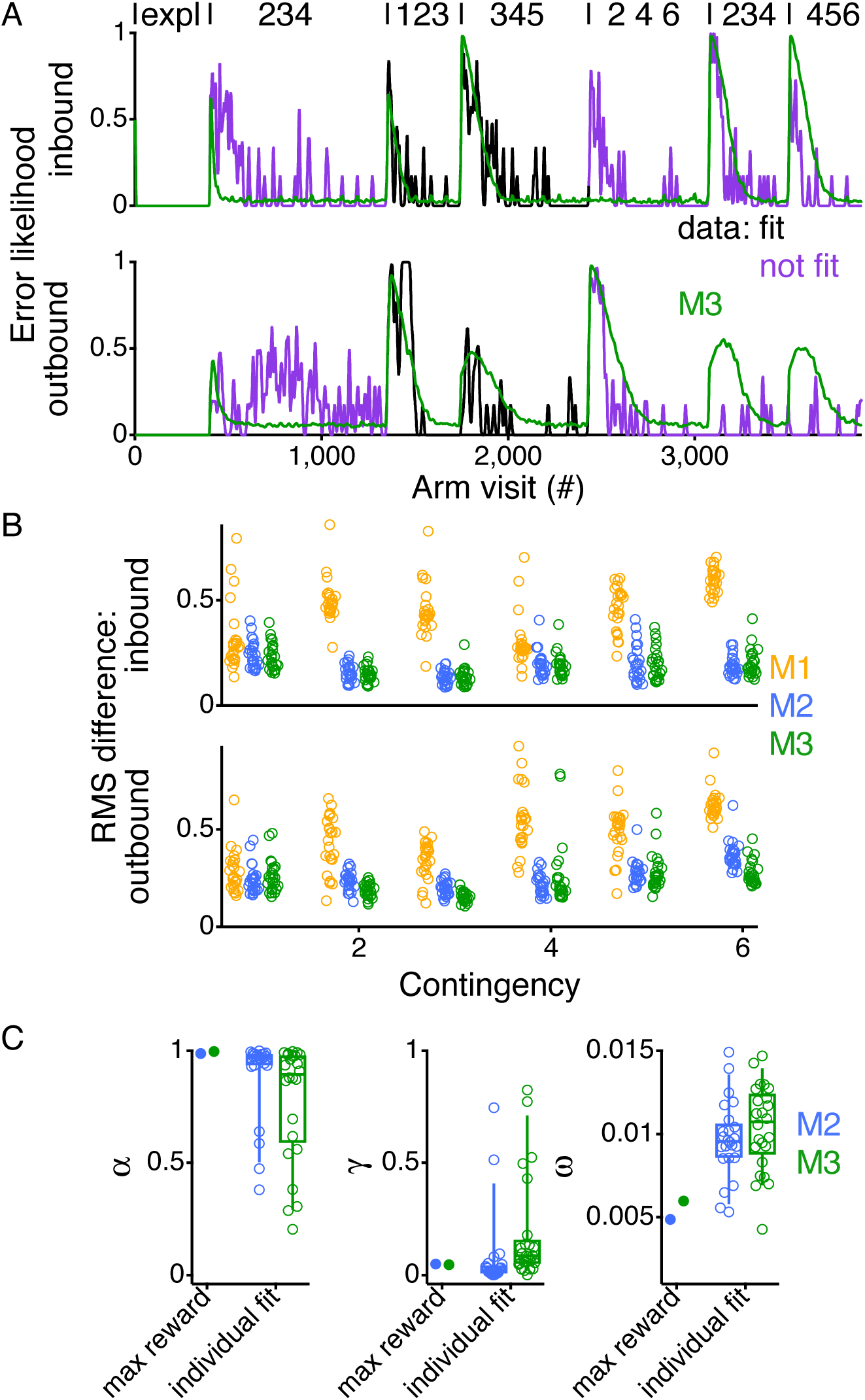
Fitting model to individual animals to capture variability between rats. (A) Inbound and outbound error likelihood for an individual animal across all contingencies (purple/black). Values smoothed with a Gaussian filter with a standard deviation of 2.25 errors and then interpolated to reflect arm visits. In green is the average behavior of 200 repeats of the model using the parameters that minimize the rms difference between the model and the animal during the second and third alternation contingencies (black). Purple indicates data that was not included in fitting the model. Contingencies indicated as in Fig 1F. (B) RMS difference between the model and the data for all animals (n = 24) for the inbound and outbound errors for each contingency for the different models. (C) Comparison of the parameters for the fits of individual animals (open circles) to the parameters that maximize rewards (closed circles) from Fig 1C. Box plots show the median, interquartile range and the range between the 9^th^ and 91^st^ percentile of the data.

### Individual model fits capture variability in behavior

M3 yielded parameter estimates for the learning-related variables that were much more variable across animals, suggesting that it might capture individual differences. When compared to M2, the M3 fits to all 24 rats had an interquartile range 7.8 times larger for *α* (0.39 vs. 0.05) (*p* = 0.0021, paired permutation test), 3.0 times larger for *γ* (0.11 vs. 0.04) (*p* = 0.0007, paired permutation test), and 1.8 times larger for *ω* (0.004 vs. 0.002) (p = 0.023, paired permutation test).

Indeed, M3 fit not only the overall structure of the learning of all animals, but also captured information about the individual learning rates of each animal. Individual animals had different overall reward rates (Fig. 1F), and thus to be able to capture differences between the animals, the model needs to show differences in reward rates between the fits of the animals.

This was the case: the fits of M3 to the individual animals correspond to different total reward rates of the model (Fig. 4A). This means that M3 provides a good substrate to explain the different performance of the rats on the task.

**Figure 4.**
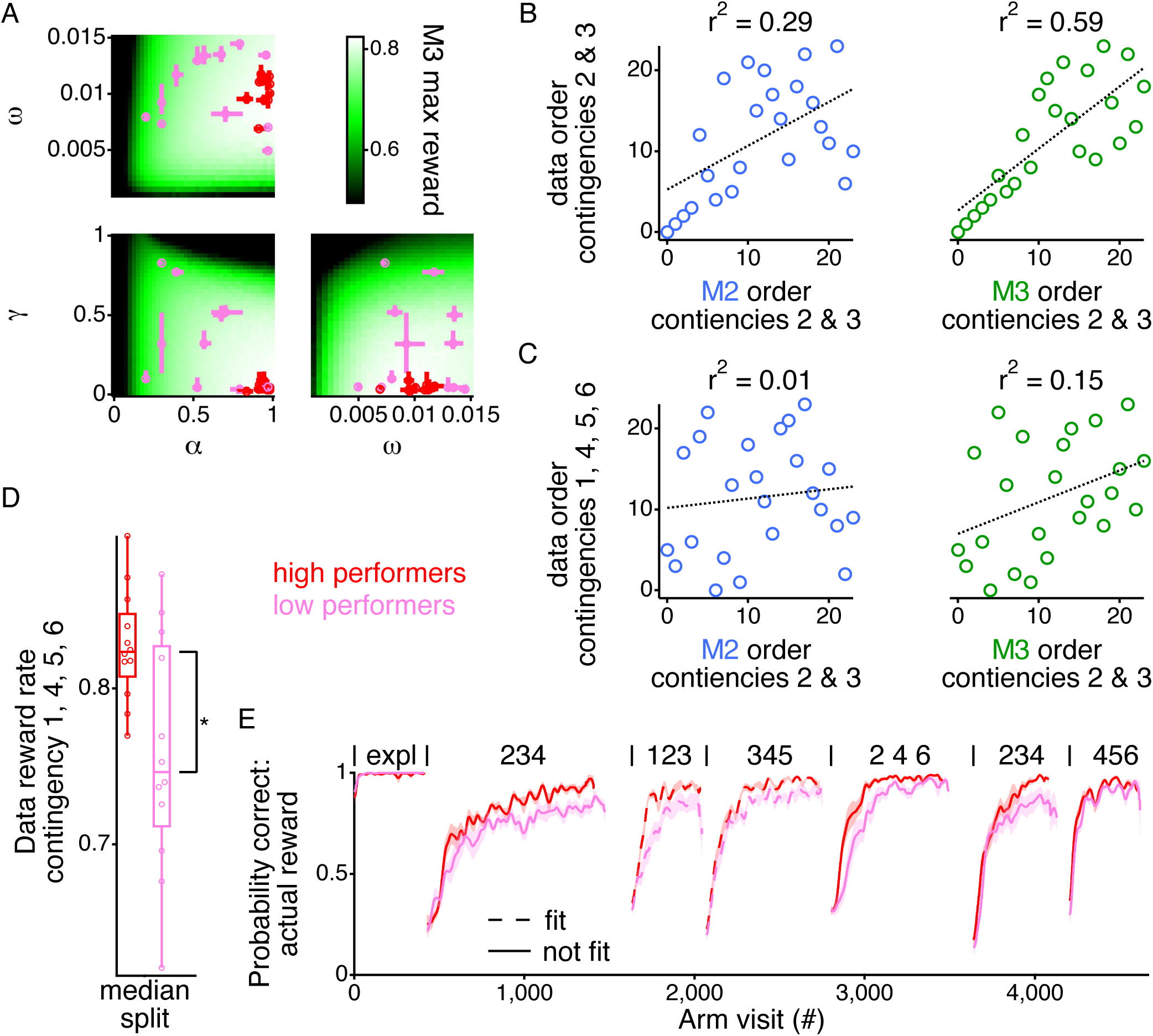
M3 captures differences in animal performance. (A) Three-dimensional space of parameters projected down on all pairs of parameters. The median and interquartile range of 20 fits for each animal is plotted as the red or pink dot with errors bars in both axes. Red and pink colors indicate the median split of animals as shown in panels D and E. Color scale in background is the maximal reward rate during the second and third contingency for the pair of parameters. For instance, for the *α*/*γ* plot, each color indicates the maximal reward value that can be obtained for the pairing of *α* and *γ*, which is found by scanning through all values of *ω*. (B) Ordering of the animals based on the actual reward rate during contingencies 2 and 3 as a function of the ordering of the animals based upon the model reward rate during contingencies 2 and 3, for M2 (left) and M3 (right). Dotted lines show a linear fit to the correlation. (C) Ordering of the animals based on the actual reward rate during contingencies 1, 4, 5, and 6 (those not fit by the model) as a function of the ordering of the animals based upon the model reward rate during contingencies 2 and 3, for M2 (left) and M3 (right). Dotted lines show a linear fit to the correlation. (F) Box plots showing the data, median, interquartile range, and 9^th^ to 91^st^ percentile for the actual reward rate of the animals during contingencies 1, 4, 5, and 6 when split by the model reward rate during contingencies 2 and 3. * indicates *p* < 0.05. (E) Average (± sem) probability correct across all contingencies for the grouping by the median split of the M2 reward rate for contingencies 2 and 3. Contingencies indicated as in Fig 1F. Solids lines indicate contingencies that were not fit by the model, and dotted lines indicate those contingencies that were fit by the model.

M3 also captures the relative performance of the animals better than M2. We ordered the animals based upon the actual reward rate the animals received during the second and third alternation contingencies and compared that to the order of the animals based on the reward rate the model received when fit with either M2 or M3. M3 better accounted for the variability in reward rates than M2 for contingences which the model was fit (Fig. 4B). M3 captured 58.8% of the variance in the ordering of reward rates of the animals during contingency 2 and 3, which was substantially larger than the 29.1% captured by M2 (*p* = 0.017, paired permutation test).

That strong correlation is a necessary, but not sufficient condition for M3 being considered a good model. A good model should also make accurate predictions on new data. We therefore asked if M3 could capture the variability in the performance of the animals for the contingencies on which the model was not fit and thereby capture something about the learning for each animal. We found that the ordering of the model reward rates from contingency 2 and 3 captured 15.3% of the variance in the performance of the animals to the contingencies that were not fit by the model (1, 4, 5, and 6), which is substantially larger than the 1.3% captured by M2 (*p* = 0.017, paired permutation test) (Fig. 4C). Importantly, M3 captured the same amount of variance in the reward rate of the animals to the contingencies that were not fit as the actual reward rates of the animals in contingencies 2 and 3 (*r*^2^ = 10.6%; *p* = 0.30, paired permutation test). That indicates that M3 does at least as good a job of predicting the reward rate of the animals as the reward rate of the animals themselves.

An examination of the overall reward rates themselves confirmed these conclusions. We performed a median split based off of the reward rate of the model to contingency 2 and 3. The higher performing half of the animals showed a significantly greater overall reward rate on the remaining contingencies (1, 4, 5 and 6) compared to the lower performing half of the animals (Fig. 4D, *p* = 0.02, rank sum test). The average performance of the higher performing half of the animals was consistently larger than the lower performing half of the animals across all of the contingencies, even though the median split was made off of the reward of the model for contingency 2 and 3 (Fig. 4E).

### Model agnostic analysis confirms importance of neighbor preference

The modeling provides strong support for the importance of the dynamic preferences for the rapid learning of this family of spatial alternation tasks. In particular, adding in the neighbor arm preference was critical for capturing the individual variability among rats in learning this task (Fig. 3C, 4). That observation led us to ask whether the neighbor bias could also account for other aspects of behavioral performance.

Consistent with this possibility, we found evidence that the neighbor bias from the exploratory period of the task relates to overall performance on the alternation task. During the exploratory period, we calculated the frequency with which each animal visits the neighboring arm. There was a range of preferences across the rats for neighboring arms during the exploratory period, which correlated surprisingly well with the average reward rate across all contingencies for each animal (Fig. 5A, *p* = 0.0016, *r*^2^ = 0.37). Thus, animals that demonstrate a stronger preference for visiting neighboring arms tend to obtain more rewards, possibly because the structure of the task includes neighboring arm visits.

**Figure 5.**
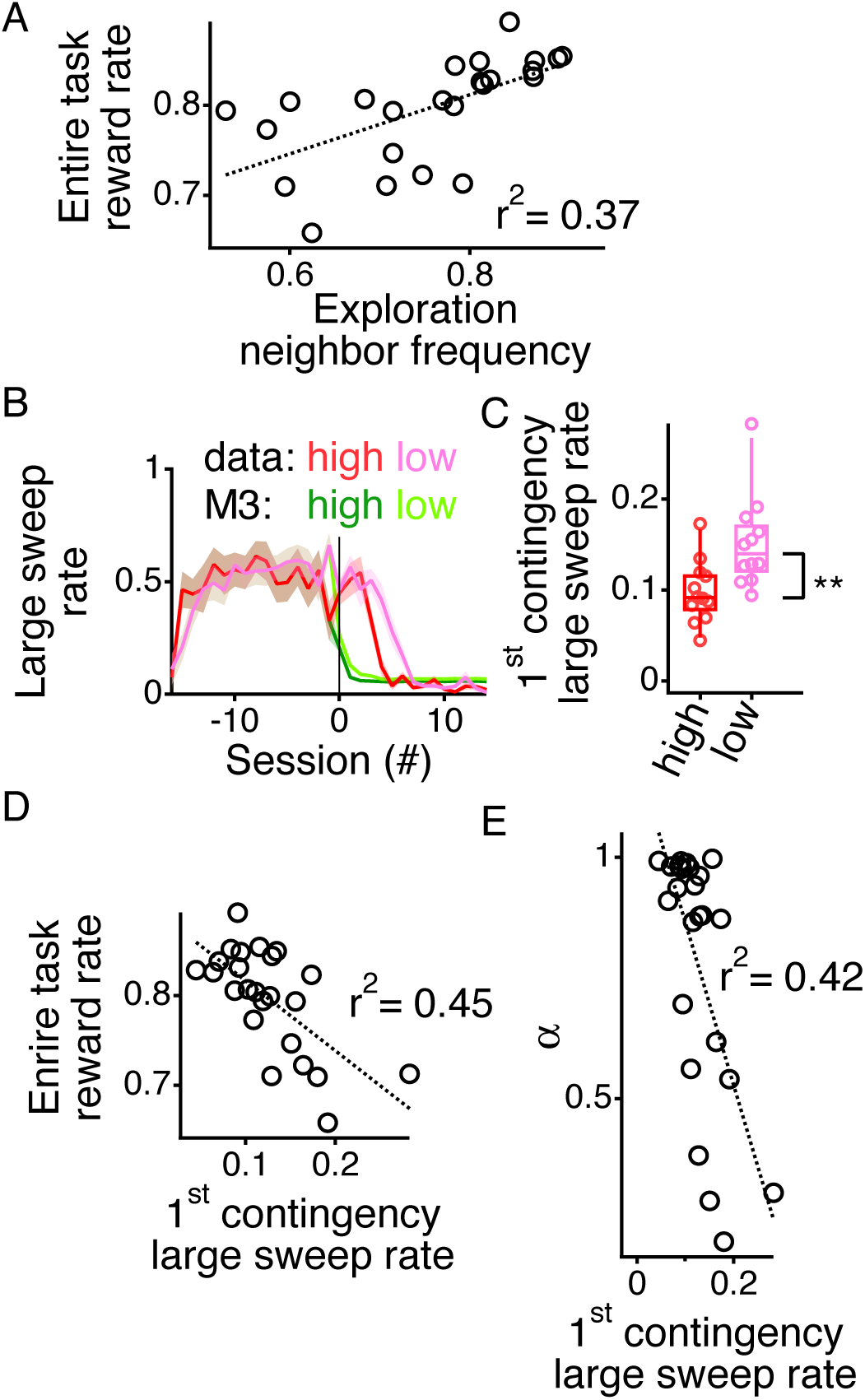
Dynamic preferences account for variability in reward rate across animals. (A) The average reward rate across all six alternation contingencies plotted relative to the average neighbor transition frequency during the exploratory period for each animal. Dotted line shows linear fit. (B) Average (± sd) large sweep rate (>3 arms) for each session during the exploratory period and first alternation contingency. Animals split into high (red) and low (pink) performers based upon median split from Fig 4. Same measurement calculated off of the 200 repeats of the model using the fitting parameters for each of the animas. M3 split based upon the same grouping as the animals. Solid vertical line demarcates the transition between exploration and the first alternation contingency. (C) Box plot showing the data, median, interquartile range, and 9^th^ to 91^st^ percentile of the large sweep rates across the entire first contingency for the high and low performing animals. ** indicates *p* < 0.005. (D) The average reward rate across all six alternation contingencies plotted relative to the large sweep rate during the first alternation contingency for each animal. Dotted line shows linear fit. (E) Learning rate (*α*) from the individual fits of model M3 to each animal (fit for 2^nd^ and 3^rd^ alternation contingency) plotted relative to the large sweep rate during the first alternation contingency for each animal. Dotted line shows linear fit.

### Additional preference governs slower learning of first alternation contingency

It is possible that learning the alternation task draws upon preferences that were not included in the model. As shown in Figure 1D&E, animals exhibit directional inertia during the exploration period. M3 did not include this preference, allowing us to ask whether any of the discrepancies between M3 and the behavior of the rats could be due to the absence of directional inertia in the model. Directional inertia leads to large sweeps across the track (Fig. 1D) and sweeps larger than 3 arms are counterproductive for the alternation task.

We asked whether there was strong evidence for including directional inertia in the model by asking if the preference to perform large sweeps during the exploratory period predicted the average reward rate throughout the task, as occurred for the neighbor transition preference (Fig. 5A). This was not the case: even though directional inertia was prevalent for the animals (Fig. 1E), there was no significant correlation between the large sweep rate (sweeps >3 arms) during the exploratory period and the total reward rate during the alternation task (Fig. S5A) (*p* = 0.4).

There was, however, clear evidence of slower learning by the rats on the first contingency, as compared to M3. This slower learning occurred in part because for the rats, but not the model, large sweeps often persist into the first contingency. We calculated the proportion of arm visits that were a part of a large sweep (>3 arms) during the exploratory period and into the first alternation contingency (Fig. 5B). The values are identical between the animals and the models fit to those animals during the exploratory period because we force each model to follow the same series of arm visits as the individual rats (see Methods). At the transition to the first contingency, M3 drops to a low baseline level of large sweeps. In contrast, the rats persist with an elevated large sweep rate after the transition to the first alternation contingency (Fig. 5B).

To provide further evidence that persistent large sweeps lead to slower learning of the first contingency, we evaluated the large sweep rates of the higher and lower performing rats, as determined by the median split from the model fit to the second and third contingency (Fig. 4C). The higher performing animals dropped their large sweep rate faster than the lower performing animals (Fig. 5B), with the higher performing rats having a lower overall large sweep rate in the first contingency compared to the lower performing rats (Fig. 5C) (*p* = 0.003, rank sum test). These two groups of animals did not show any difference in large sweep rates during the exploratory period (*p* = 0.55, rank sum test). This is consistent with the higher performing animals more quickly learning to not perform large sweeps.

If so, then animals that learn faster should be able to more quickly overcome their preference for directional inertia. Indeed, that was the case. We calculated the large sweep rate from the first contingency, where fewer large sweeps would be expected to be associated with faster suppression of this preference. We found a strong inverse correlation between the reward rate for the entire task and the first contingency large sweep rate (Fig. 5D) (*p* = 3.3 × 10^-4^, *r*^2^ = 0.45). Consistent with the removal of the large sweeps being a function of the learning capacity of the animals, there was also strong inverse correlation between the learning rate, *α*, of the model (fit only to the second the second and third contingency) and the first contingency large sweep rate (Fig. 5E) (*p* = 6.0 × 10^-4^, *r*^2^ = 0.42), with *α* also accounting for a large fraction of the variance of the overall reward rate (*p* = 1.2 × 10^-4^, *r*^2^ = 0.38).

Finally, we asked whether the large sweep rate, and by extension the learning rate, capture a different aspect of the reward rate variability than that which is correlated with the neighbor transition frequency during the exploratory period (Fig. 5A). We found that it does: the neighbor transition frequency during exploration did not correlate with the large sweep rate during the first alteration contingency (Fig. S5B) (*p* = 0.2).

In combination, the neighbor transition frequency during the exploratory period and the large sweep rate during the first alternation contingency account for 64.6% of the variance in the reward rates of the animals across the entire alternation task. We calculated the overall variance explained by fitting a multifactorial linear regression relating the transition frequency and large sweep rate to the overall reward rate during the alternation task. Consistent with the large sweep rate being correlated with the learning rate of the animals, the neighbor transition frequency during the exploratory period and the learning rate of the model (fit only to the second and third contingency) account for 58.2% of the variance in the reward rates of the animals across the entire alternation task. The slight increase in variance captured by including the large sweep rate in the first contingency over including the learning rate of the model occurs due to the fact that the large sweep rate in the first contingency is directly related to the amount of reward in the first contingency. If we compare the reward rate for contingencies 2 -6 the neighbor transition frequency combined with the large sweep rate accounts for 64.9% of the variance; whereas the neighbor transition frequency combined with the learning rate accounts for 66.4% of the variance.

Taken together, these findings indicate that the lack of directional inertia contributed to the faster learning of the first alternation contingency by M3 as compared to the rats. But, after the transition from exploration to the first contingency, the variability of the animals with respect to directional inertia was well captured by the variability in the associated model learning rates.

### Model accurately predicts animal behavior

As an additional test of the model and the importance of combining dynamic preferences and memory to describe the way the animals learn, we examine the ability of the model to predict the course of learning on the contingencies on which the model was not fit (1, 4, 5, and 6). We first define a metric for model accuracy. Each time the model is run, it generates a set of choices and a corresponding set of inbound and outbound errors. Up until this point, we combine 200 repeats of the model to define the average inbound and outbound error rates. Here we assess the variability of the model by measuring the rms difference between the inbound and outbound errors of an individual run of the model and the average inbound and outbound error rates of the model. If the model is a good fit for an animal, then the errors of the animal should be similar to those individual runs of the model (Fig. 6A, S6), and thus should have a similar rms difference to that of individual runs of the model to the average error rates from the 200 repeats. Model accuracy was therefore defined as the proportion of the individual model runs with larger rms differences than the data, and thus values less than 0.05 indicate significant deviations of data from model at *p* < 0.05, while large values indicate non-significant deviations.

**Figure 6.**
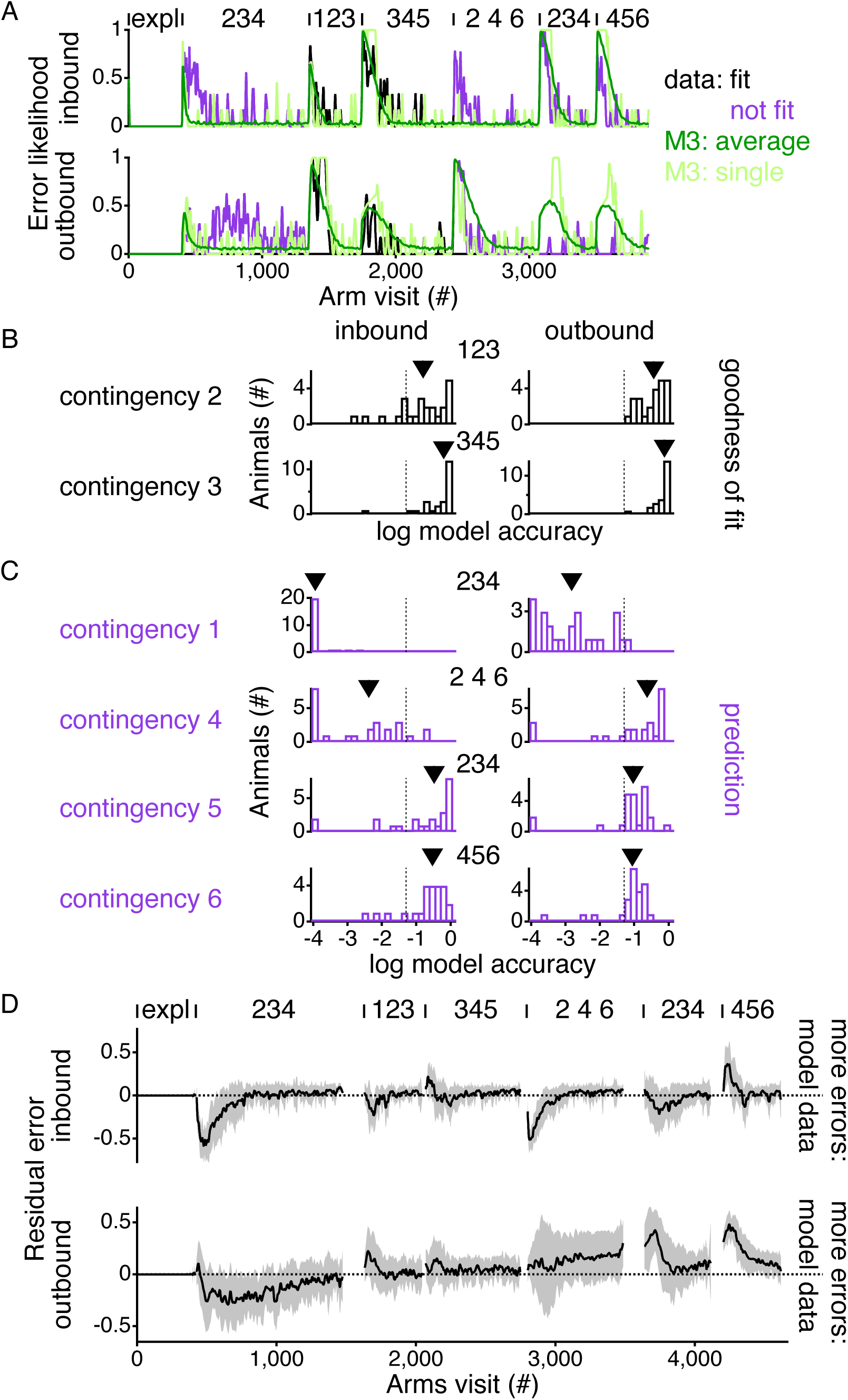
Model predicts performance of animals on contingencies that were not fit. (A) Inbound and outbound error likelihood for an individual animal across all contingencies (purple/black). Values smoothed with a Gaussian filter with a standard deviation of 2.25 errors and then interpolated to reflect arm visits. In dark green is the average behavior of 200 repeats of the model using the parameters that minimize the rms difference between the model and the animal during the second and third alternation contingencies (black). Purple indicates data that was not included in fitting the model. In light green is the inbound and outbound error likelihood for a single repeat of the model using the parameters from the fit to the individual animal. (B) Histograms across animals of the log of the model accuracy of M3 for the two contingencies (2&3) that were used to fit the model. (C) Histograms across animals of the log of the model accuracy of M3 for the contingencies that were not used to fit the model. For B&C above each pair of histograms are the arm numbers that were rewarded during that contingency. Model accuracy is the probability of individual runs of the model providing an error greater than the error of the data (Fig S6). Vertical dotted lines show *p* = 0.05 for comparison. Arrowheads point to value of the median for the model accuracies. (D) Difference between the error likelihood for the rats and the model fit to the individual rats, averaged across all rats (± standard deviation). Positive residual values indicate that the model had higher error likelihoods and negative residual values indicate that the model had lower error likelihoods. For panels A&D contingencies indicated as in Fig 1F.

The model accuracy confirms the goodness of fit of the model to contingencies two and three, the contingencies to which it was fit (Fig. 6B). The majority of the animals show model accuracies far larger than 0.05 both for the inbound and outbound errors (table 1), creating a median *p* value for the population greater than 0.05, which indicates that the model fits of the error rates of the animals during contingency two and three cannot be distinguished from individual runs of the model.

**Table 1.** Model accuracy statistics. Fraction of animals whose inbound and outbound errors cannot be distinguished (*p* > 0.05) from the model as well the population median p-value. Contingencies 2 &3 were fit to the model, the remaining contingencies were not included in the fit and therefore function as predictions of the model.

| contingency | inbound |  | outbound |  |
| --- | --- | --- | --- | --- |
| | fraction of animals $p > 0.05$ | population median p-value | fraction of animals $p > 0.05$ | population median p-value |
| 1 | 0/24 | 0.001 | 1/24 | 0.002 |
| 2 | 17/24 | 0.16 | 24/24 | 0.36 |
| 3 | 23/24 | 0.62 | 24/24 | 0.75 |
| 4 | 3/24 | 0.004 | 18/24 | 0.24 |
| 5 | 18/24 | 0.33 | 20/24 | 0.09 |
| 6 | 20/24 | 0.29 | 20/24 | 0.10 |

The models also provide good predictions for many of the remaining, nonfit, contingencies. Outbound errors on contingencies 4, 5, and 6 were well described by the model, as were the inbound errors on contingencies 5 and 6 (Fig. 6C, table 1). Of these the most surprising is that M3 fit to the second and third contingency predicted the outbound errors of the fourth contingency (Fig. 6C). This contingency, 2-4-6, is by far the hardest contingency, as it forces the animals to continue to alternate arms but to do so whilst skipping neighboring arms. Not surprisingly, the fits to the first contingency are poor, due at least in part to the initial directional inertia of the rats.

### Deviations of the behavior from the model point to generalization about the task

Even though M3 fit to the second and third contingencies did a good job of predicting the outbound errors of the subsequent and unfit contingencies, we were surprised that the outbound errors of contingency five and six were less well fit than the outbound errors of the fourth, and hardest contingency. We quantified this and found that the distribution of model accuracies shifts is indeed worse for the outbound errors of the fifth and sixth contingencies, where the median of the distribution is 0.092 and 0.099, respectively (Fig. 6C). These accuracies are far worse than the outbound fit accuracies for the second (0.363; *p* = 1.8 × 10^−4^+ and *p* < 1 × 10^−5^ respectively, paired permutation test) and third (0.752; *p* < 1 × 10^−5^ and *p* < 1 × 10^−5^ respectively, paired permutation test) contingencies, and, surprisingly, the medians are also worse than the outbound fit accuracy for the fourth contingency (0.237; *p* = 3.8 × 10^−3^ and *p* = 2.5 × 10^−4^ respectively, paired permutation test), even though the fifth and sixth contingencies are of the standard variety with neighboring arms being the outer arm.

We then asked whether these less accurate fits were associated with better or worse performance by the model: if the model performs worse than the animals it would indicate more rapid learning and lower error rates in the animals that could result from generalization where the animals had extracted information about the structure of the task. To explore that possibility, we calculated the average difference between the inbound and outbound errors of the individual rats and the inbound and outbound errors of the model fits (Fig. 6D). Consistent with the possibility of generalization, the model makes more outbound errors at the beginnings of the fifth and sixth contingencies. This is in contrast to the performance on early contingencies (1 and 4) where the model tended to make fewer errors.

## Discussion

Can we rely on our intuition to understand behavioral performance in complex tasks? In the context of spatial alternation, differences in learning rates are typically interpreted as reflecting differences in the quality of each animal’s memory for past experiences^12, 13,15’17^’^33^. We developed an automated six arm spatial task and exposed rats to both an initial exploration period and a series of alteration contingencies where the animal had to alternate among different subsets of arms (Fig. 1). We then developed a series of RL models, first using memory alone and then, when that model proved insufficient, incorporating specific dynamic preferences that reflect favored arms or favored transitions between arms (Fig. 2).

As we also found for the simpler, three-arm task^18^, the incorporation of these dynamic preferences was sufficient to produce a model that can learn the spatial alternation task as rapidly as the rats (Fig. 2&3). That model also identified different learning parameters across animals and was able to predict individual animal behavior on data to which it had not been fit (Fig. 4&6). The specific preferences added to fit the data included a neighbor transition preference that could be estimated from the initial exploration period. The strength of that preference for individual animals was highly predictive of the total amount of reward they received throughout the task (Fig. 5A). Our results demonstrate that the dynamics of learning can be captured with relatively simple models that combine memory with dynamic preferences.

### Importance of assessing preferences before task learning

Traditionally, when studying spatial alternation behavior, animals experience the space of the task at the same time that they experience the rules of the task^18^. This conflates the learning of space with the learning of the task. Furthermore, it prevents any direct and independent measurement of the preferences that the animals bring to the task.

Here we have separated these two aspects of the behavior. In so doing we directly measured the preferences that rats have in exploring the six arms of the track (Fig. 1 B-E). The presence of these preferences indicates that they could be utilized while learning the task. We have presented model dependent (Fig. 4&6) and model agnostic (Fig. 5) evidence that rats dynamically utilize these preferences to enable their rapid learning of the spatial alternation task.

### Utilizing individual variability for model refinement

The procedures of model generation, testing and refinement are exactly those of formulating and falsifying hypotheses about possible underlying causes of the observed behavior. Such hypotheses are typically componential—involving different potential processes (such as learning rules) and parameters (such as subject-specific learning rates or weights governing the impact of alternative mechanisms). Testing modelling hypotheses of this sort requires determining the values of the parameters that fit the data, and balancing the quality of the fits against the complexity of the models^42^. New and additional hypotheses then arise from features in the residuals of the fits to the data.

Since the modelling procedures can sometimes be buried or even ignored, we took the opportunity to lay out the logic of their development and motivate their form through the intricacies of the data. In doing this, we have presented a series of related models that increasingly capture the behavior of the animals across multiple stages of the task.

Through refinement, we developed an RL agent that could perform the task as quickly as the rats and fit that model to capture the individual behavior. We laid bare the logic of the changes that we made to the RL agent to develop the final model. As the alternation task cannot be described as a Markov decision process, a working memory RL agent (M1)^35^ could learn the task, but did so far slower than the animals for all but the first alternation (Fig. 2). Incorporating dynamic preferences, motivated by the exploratory behavior of the rats, into the RL agent (ultimately model M3) enabled it to learn as rapidly as the rats (Fig. 2). Note that all models (M1 -M3) can be described as model-free agents with temporally-sophisticated representations; thus, our result confirms Akam et al.’s^43^ observation, that one need not necessarily appeal to more computationally sophisticated, model-based, components^28^ to account for all aspects of fast learning, but rather take appropriate account of structural contributions associated with preferences.

Since, by their very nature, models offer incomplete and simplified representations of phenomena of interest, it is critical to be able to define how to decide where to stop the formal modeling process. Through the various model iterations, we had to decide both when to continue refining the model, and when to stop doing to so. The initial drive to continue the modeling was obvious. M1 couldn’t learn the task nearly as rapidly as the animals (Fig. 2). However, with M2, the need to continue was far more subtle. M2, largely, had the capacity to learn the task as rapidly as the rats; however, when we fit M2 to the individual animals the parameters mostly clustered around the parameters that maximized the amount of reward that M2 could receive (Fig. 3C). This indicated that there was still something fundamentally missing in the agent. None of the subtlety and richness of the individual variability between animals clearly apparent in Fig. 1F&G existed within the parameters for the fits of M2. Finally, with M3, the parameters from the fits to the individual animals varied away from the parameters that maximized reward for that agent. This indicated that we could consider no longer continuing the process of improving the modeling.

The decision to stop the modeling at that point rested upon somewhat different factors. Even though M3 fails to capture certain identifiable aspects of the behavior of the rats, it closely matches different features of the way in which the individual animals learned the task (Fig. 6), and predicted the learning to contingencies that were not fit by the model (Fig. 4). It was the combination of capturing variability across animals and predicting features of the behavior outside of the fit that suggested that we might stop refining the model further. To go beyond this, it would be desirable to refine the paradigm, for instance to put the ‘skip’ contingency (2-4-6) at different points to examine its role in generalization; or to systematize the length of engagement with each contingency to study the possible decrease in learning rate as performance improved.

### Model successes and limitations point to continual learning and generalization

Animals seamlessly learn tasks over many different timescales, a characteristic with which machine learning and artificial intelligence are just starting to grapple^44, 45^. Such continual learning can be precisely defined using our presented modeling approach. Our model, fit to just two of the contingencies, predicts the behavior of the animals in other contingencies (Fig. 6). That indicates that there need be no new type of learning for the fit and predicted contingencies, even though the specific application of the rule changes. However, in places where the model does not predict the behavior, that provides specific times where the animal could change its learning and provide experimental substrate to better understand where and how animals continually learn.

The places where the model less well predicts the behavior of the animals also allows for the generation of specific hypothesis as to what the animals might be doing. The model made more outbound errors on the final alternation contingencies and fewer inbound errors of the fourth contingency (Fig. 6). We speculate that it is here that M3 is compromised by its inherent model-free nature and that the rats deviated from the RL agent because the rats have generalized by learning the structure of the task, enabling them to use that information to speed up the learning of a given contingency (or possibly slowing learning when their expectations fail to match reality). It will be interesting to study the timing and nature of this potential generalization further. Does the generalization only happen due to the number of contingencies, or is there something about the animals’ experience of the skip arm contingency that allows them to generalize the alternating nature of the task more competently?

Finally, using an extended and changing task combined with the modeling allowed us to find systematic differences between animals in the way that they learned the task (Fig. 4). These results are consistent with mice studies showing consistent learning capacities across tasks^46, 47^. Furthermore, the link between exploratory behavior and learning ability (Fig. 5) is also consistent with previous mouse studies^48^. This sets the state for future studies that would identify individual differences in the circuitry supports the learning of these behaviors.

## Methods

### Animals

All experiments were conducted in accordance with University of California San Francisco Institutional Animal Care and Use Committee and US National Institutes of Health guidelines. Rat datasets were collected from Long Evans rats, ordered from Charles River Laboratories, that were fed standard rat chow (LabDiet 5001). To motivate the rats to perform the task, reward was sweetened evaporated milk, and the rats were food restricted to ∼85% of their basal body weight.

Two cohorts of rats, comprised of 6 males and 6 females each, were run on the automated behavior system. There were no systematic differences in reward probabilities between the male and female rats within the two cohorts (Fig. S1C), so data from all animals were aggregated for subsequent analyses. The entire behavior took place over the course of 22 days for the first cohort and 21 days for the second cohort. The first cohort ran an extra day on the initial exploratory behavior, where the animals received rewards after visiting any arm of the track. At the start of the behavior the first cohort of rats were 4 -5 months old, and the second cohort of rats were 3 -4 months old.

### Automated behavioral system

The automated behavior system was custom designed and constructed out of acrylic. All parts of the behavior system were enclosed with walls. There were different symbols on each arm of the track serving as proximal cues, and there were distal cues distinguishing the different walls of the room. Pneumatic pistons (Clippard) opened and closed the doors. Python scripts, run through Trodes (Spike Gadgets), controlled the logic of the automated system. The reward wells contained an infrared beam adjacent to the reward spigot (Fig. S1A). The automated system used the breakage of that infrared beam to progress through the logic of the behavior. In addition to the infrared beam and the spigot to deliver the reward, each reward well had an associated white light LED (Fig. S1A).

The sequence of operations of the track for the set of behaviors are: 1) the doors open to clear the path from a single rest box to the track. Concurrently, the lights linked to all of the reward wells on the track turn on (Fig. S1A). 2) On the first break of a track reward well beam (Fig. S1A) following the opening of the doors, the door to the track closes, thus starting the session of that animal. The animal then has a fixed maximum number of trials for its session, and the session ends when either that maximum has been reached or following a time limit of 30 minutes (see Methods). Only one animal ever reached the time limit (see Methods). 3) Upon breaking the beam at the reward well at the last trial of the session, all of the reward well lights turn off, and the doors reopen to allow for passage back to the appropriate rest box. Concurrently, the light to the reward well in that rest box turns on. 4) Upon breaking the beam of the rest box reward well, the doors to the track close and the well delivers reward. The light of the rest box reward well turns off after reward delivery. 5) The doors to the track for the rest box for the next subject open, and the process repeats itself.

Each cohort of rats were divided into groups of four animals. The same groups were maintained throughout the duration of the experiment. Within a group, a given rat was always placed in the same rest box, and the four rats of a group serially performed the behavior. The rats had multiple sessions on the track each day. During the exploratory period of the behavior, the duration of a session was defined by a fixed number of rewards. During the alternation task the duration of a session was defined either by a fixed number of center arm visits and at least one subsequent visit to any other arm, or a fixed amount of time on the track (30 minutes), whichever came first. During the alternation contingencies there were 3 sessions each day. For the first day of the first alternation contingency there were 10 center arm visits per session, for the second day of the first contingency and the first day of all other contingencies there were 20 center arm visits per session, and for all other days there were 40 center arm visits per session. Only one of the female rats reached the time limit, and it did so for only two sessions toward the beginning of the first alternation contingency. For that one female we incorporated the trials that she ran on those sessions and did not distinguish the time out sessions for the analyses.

The algorithm underlying the exploratory part of the behavior had only one rule. Reward was delivered for any infrared well beam break if and only if the current well infrared beam break was immediately preceded by an infrared beam break at any other well. This prevented the animals from getting continuous reward at a single arm, and ensured the rats visited at least two of the arms.

The algorithm underlying the spatial alternation task was such that three arms on the track had the potential for reward within a given contingency, for example during the contingency at arms 2-3-4, arms 2, 3, and 4 had the potential to be rewarded, and arms 1, 5, and 6 did not. Of those three arms we will refer to the middle of the three arms as the center arm (arm 3 in the above example) and the other two arms as the outer arms (arms 2 and 4 in the above example). Reward was delivered at the center arms if and only if: 1) the immediately preceding arm whose reward well infrared beam was broken was not the center arm. Reward was delivered at the outer two arms if and only if: 1) the immediately preceding arm whose reward well infrared beam was broken was the center arm, and 2) prior to breaking the infrared beam at the center arm, the most recently broken outer arm infrared beam was not the currently broken outer arm infrared beam. The one exception to the outer arm rules was at the beginning of a session, following the first infrared beam break at the center arm, where only the first condition had to be met, if neither of the outer arms had yet to be visited.

For the running of the behavior, the infrared beam break determined an arm visit (Fig. S1A); however, the rats would sometimes go down an arm, get very close to the reward wells, but not break the infrared beam. Therefore, for all of the analyses described, an arm visit was defined as when a rat got close to a reward well. The times were extracted from a video recording of the behavior. These missed pokes were more frequent at the beginning of a contingency (Fig. S1D), but overall were not that common. This proximity-based definition of an arm visit added additional arm visits to those defined by the infrared beam breaks, and by definition none of them could ever be rewarded, nor alter the logic of the underlying algorithm. However, because of the non-Markovian nature of the reward contingency, the missed pokes could affect the rewards provided for subsequent choices.

The different spatial alternation contingencies (Fig. 1F) were chosen to present increasing challenges and multiple learning opportunities. The transition from the first (2-3-4) to the second (1-2-3) contingency was designed to be relatively easy, since performing 2-3-4 would allow a rat to readily find the central arm of the new contingency. Finding this arm is critical to gaining consistent reward. The transition from the second (1-2-3) to the third (3-4-5) contingency was designed to be harder since the central arm (4) of the new contingency is not included in 1-2-3. The fourth (2-4-6) contingency was designed to be the hardest, since the animals have to skip an arm to get to the correct outer arm of the contingency. The fifth (2-3-4) and sixth (4-5-6) contingencies were chosen for comparison with the first three contingencies to understand the evolution of the ability of the animals to perform the task and generalize from previous experience.

As opposed to behaviors designed to study asymptotic performance, there need not be strict criteria on a per animal basis for switching between the contingencies since the purpose of this task was to understand the continual learning and behavior of the rats. Furthermore, the automated system matched the number of inbound rewards of the animals, for all of the animals that did not reach the time limit, ensuring that all animals had similar learning opportunities. We therefore switched to a new contingency the day after >80% of the animals received >80% reward over the course of a session. That ensured that by the time each contingency switched almost all of the rats reached at least ∼80% correct on a session during each contingency (Fig. S1B).

### RL agents

For this behavior we chose a simplified output as the modeled feature: visiting arms. The nature of the algorithm that governs the behavior led to the choice of arm visits for the model, as arms visits are the only factor taken into account when evaluating rewards.

Given that each spatial alternation task could be framed as a partially observable Markov decision process, we adapted the working memory model of Todd et al.^35^ as the basis for our series of RL agents. The models specify rules governing propensities m(a, 5) that contain the preferences of the agent of choosing arm a when the state is s. Models differ in terms of what counts as the state, and also according to the various terms whose weighted sum defines the propensity.

In the first agent (M1) the state is defined as the current arm location, *s_t_* = *a_t_*, of the agent. In all subsequent agents the state is defined as the combination of the current arm location of the agent and the immediately preceding arm location of the agent, *s_t_* = {*a_t_*_−1_, *a_t_*}. This is a simplification from the Todd et al. model, whereby *a_t_*_−1_ is always placed into the memory unit, effectively setting the gating parameter for the memory unit to always update the memory unit. Then, the first component of *m(a, s)* for all models is *b(a, s)*, which is a 6 × (6 + 1) or 6 x (36 + 6 + 1) matrix containing the transition contingencies to arm a from state *s*. The reason for the additional states beyond just the 6 arms or 6×6 arms by previous arms is to include the rest box in the possible locations to allow for the inclusion of the first arm visit of a session. In so doing that adds 1 additional state to model M1, and 6+1 additional states into the subsequent agents since the animals can be located in the rest box and can be located at any of the 6 arms having previously been in the rest box.

To provide the agents with additional spatial and transitional preferences we added components to the transition propensities. The first is an arm preference, *b*^*i*^(*a*) that is independent of the current state of the animal. The second is a preference for visiting arms that neighbor in space the current arm, 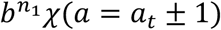, where *χ*() is the characteristic function that takes the value 1 if its argument is true (and ignoring arms outside the range 1…6) and 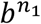 is the (plastic) weight for this component. The third is a preference for visiting arms that are two removed, in space, from the current arm, 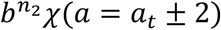. The neighbor arm preferences contain only single values, the preference to go to a neighboring arm, independent of the current arm location. The neighbor preferences were applied equally in both directions when possible (i.e. if the agent was at the end of the track the neighbor preference could only be applied to one direction).

To determine the probability of visiting each of the arms from a given state, the total propensity is passed through a softmax such that:

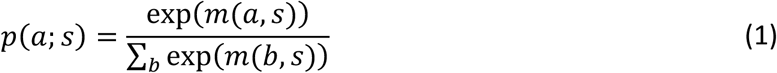

The agent’s visit is then determined by a sample from this distribution. The choice of arm then determines the reward, *r*, which is either 0 or 1, based on the algorithm that governs the spatial alternation task. The probability of revisiting the current arm is set to zero, and the probabilities of going to the remaining arms sums to 1.

The model uses the REINFORCE policy gradient method^34^ within the actor-critic framework of temporal difference learning, to update the propensities in the light of the presence or absence of reward. To do this, the agent maintains a state-long-run-value approximation, *V*(*s*), which functions as a lookup table, with one component for each state. The reward determines the state-value prediction error:

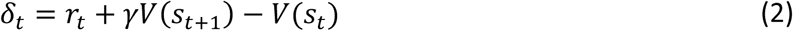

where *γϵ*[0,1] is a parameter of the model called the temporal discounting factor, which determines the contribution of future rewards to the current state.

δ*_t_* is then used to update the preferences all of the components of the propensities and *V*(*s*). The state-based transition component is updated according to the rule:

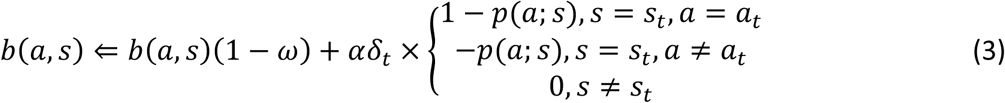

where *αϵ*[0,1] is a parameter of the model called the learning rate, which determines the amount by which all components of the propensities change based on the new information. *ωϵ*[0.001,0.015] is also a parameter of the model called the forgetting rate, and determines how the propensities decay. The independent arm preference is updated according to the rule:

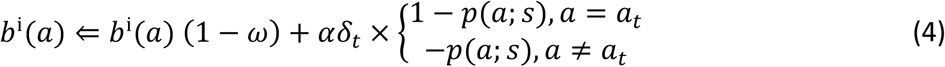

The strength of the neighbor arm preferences are updated according to the rule:

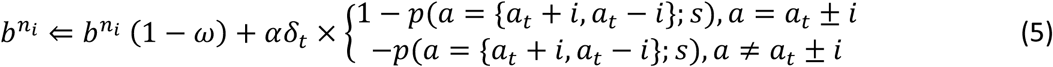

where i is either 1 or 2 depending on whether the propensity being calculated is the immediate neighbor preference or the 2 arm away preference. And, finally, the state-value approximation is updated according to the rule:

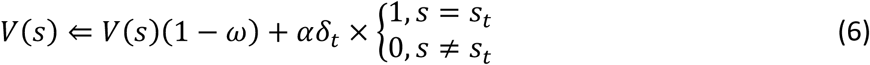

The learning, *α*, and forgetting, *ω*, rates were the same for all of the updating rules. This does not need to be the case, but since we found that a single learning and forgetting rate fit the data well, we did not feel there was a need to increase the complexity of the models by increasing the number of parameters.

### Model fitting

The model was implemented in C++ and run and fit within Igor Pro (Wavemetrics). There were 7 arms at which the agent could be located, 6 track arms and 1 rest box “arm;” whereas, there we only 6 arms to which the agent could transition. That means that the model implemented the transition from the rest box to the track but did not model the return to the rest box from the track, this was done so that all track arm visits would be included in the analyses. For the working memory version of the model, there were, therefore, 43 states in which the agent could find itself. 36 states (6^2^) for all combinations for both the previous and current arm being one of the 6 track arms (6 of the states could never be visited since a return to the same arm is not allowed), an additional 6 states for the current arm being one of the 6 track arms and the previous “arm” being the rest box, and a final 1 state for the agent starting from the rest box.

We fit the various agents to individual animals by using an Approximate Bayesian Computation method. We found the parameters that minimized the average rms difference between the inbound and outbound errors of the individual animal and of the average of 200 different repeats of the model. The inbound and outbound fitting errors were summed with equal weighting to create the final fitting error. We used simulated annealing and ran the optimization at least 4 different times from different initial conditions. We chose the parameters with the minimal error. For each run of the model we used the same random number generating seed to minimize the random fluctuations between parameter sets^42^.

We evaluated the error landscape of the fits to determine whether there were clear global minima for each animal. We found that there were indeed global minima that were distributed across the parameter space. Our fitting procedure reliably determined the vicinity of the global minima (see Fig. S4 for an example), indicating that the differences among animals are interpretable and reflect differences in behavior.

### Population statistics

For testing violations from randomness of the population, we consider a random effects model. Let *θ* be the population probability of randomness. We construct a frequentist test of the null hypothesis that *θ* = 0.5 against the one-tailed alternative that *θ* < 0.5.

If we had *m* subjects we knew were random and *n* subjects we knew were not, with *m* + *n* = *N*, then the frequentist probability associated with the null hypothesis would depend on the tail probability of the fair binomial distribution for values as, or more extreme than *n*:

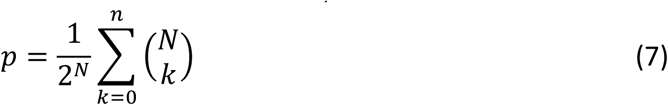

In our case, we have subject i with permutation tested probability *P*(dtat_i_|random = *ϕ*_*i*_. Thus, we have probabilities such as: 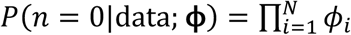, 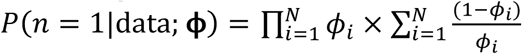 etc. Thus, we have

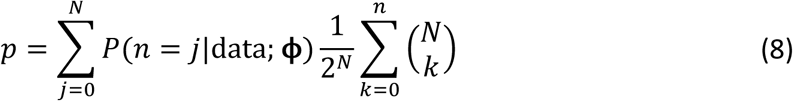

In practice, we compute this by sampling *P*(*n* = *j*|data**ϕ**). This makes the three p-values for the different exploratory preferences of the rats: 1.08e-06, 6.31e-08 and 1.75e-06, respectively for the max arm probability, neighbor transition and directional inertia.

## Data and code availability

All data and code will be made available upon reasonable request.

## Acknowledgements

We thank E. Vertes and C. Gagne for helpful discussion and A.K. Gillespie for technical assistance. This work was supported by grants from the Jane Coffin Childs Memorial Fund for Medical Research (D.B.K.), the UCSF Physician Scientist Scholars Program (D.B.K.), an NIH R25 (R25MH060482) (D.B.K), the Simons Foundation for Autism Research grant (291584) (L.M.F.), the Howard Hughes Medical Institute (L.M.F.) and the Max Planck Society and the Humboldt Foundation (P.D.).

## Author Contributions

D.B.K., L.M.F. and P.D. designed the study, D.B.K. and E.A.M. collected the data, D.B.K. and P.D. developed the models, D.B.K. and Z.Y. analyzed the data, D.B.K. designed the automated behavior system, D.K.M. and D.F.L. developed the data acquisition system, and D.B.K., L.M.F. and P.D. wrote the manuscript.

---

**Supplementary Figure 1.**
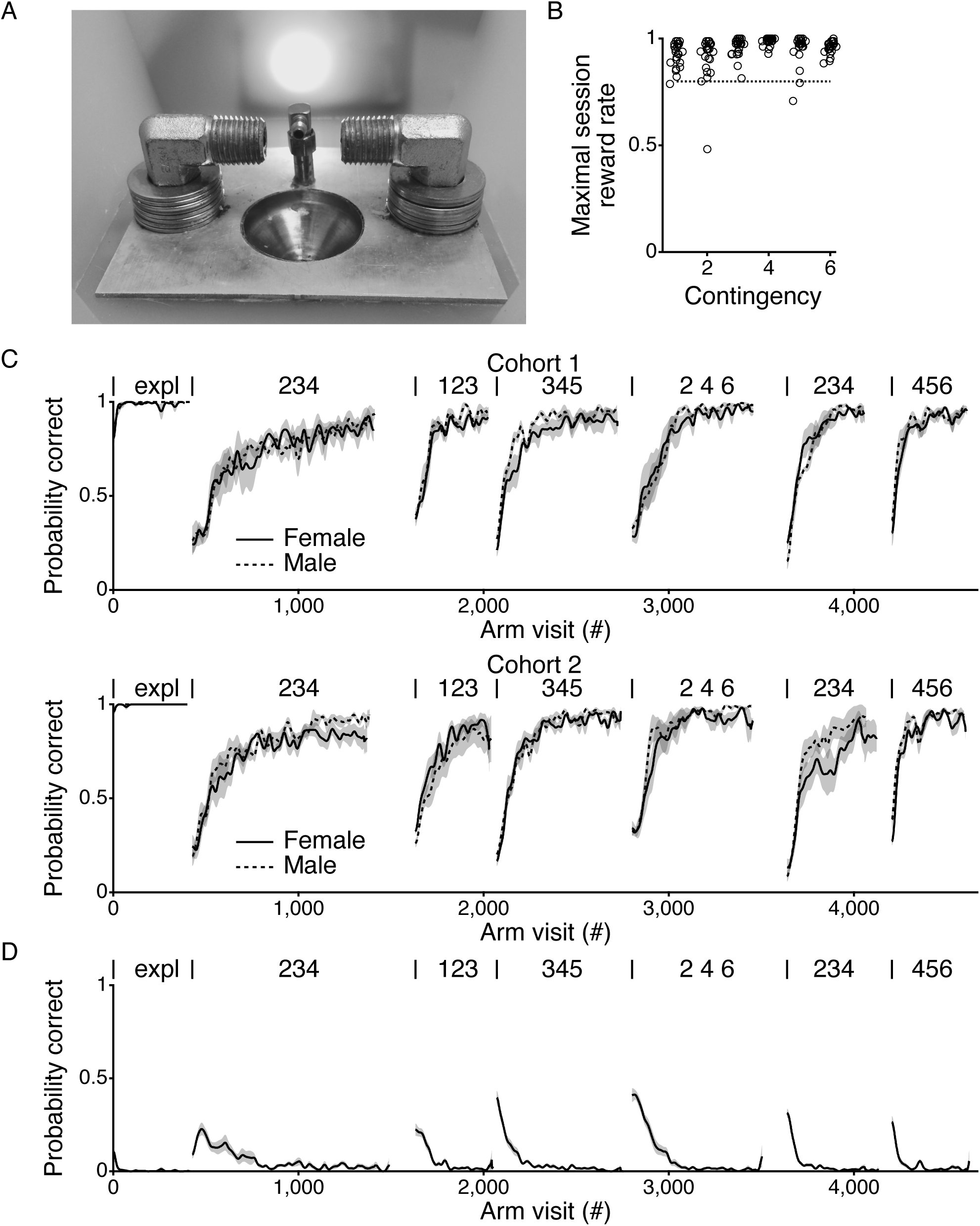
Males and females perform the behavior comparably across the two cohorts. (A) Picture of a reward well showing the spigot through which milk is delivered, flanked by an IR LED and phototransister, encased in metal elbows, to detect the position of the animal. Any unconsumed milk exited the track through the drain below the spigot. A light is illuminated directly behind the reward well when there is potential for reward delivery (see methods). Reward wells were made entirely out of metal. (B) Maximal reward rate in a session for each contingency and for all animals. Horizontal dotted line demarcates 80% correct. (C) Average probability of getting a reward for the male (dotted line) and female (solid line) rats in the first (top) and second (bottom) cohort. Within each contingency, curves smoothed with a Gaussian filter with a standard deviation of 10 arm visits and then averaged across the different animals. Thickness of the line indicates the sem. Contingencies indicated as in Fig 1F. (D) Average missed poke likelihood across all contingencies. Averaged across all rats. Thickness of line indicates sem. Contingencies indicated as in Fig 1F.

**Supplementary Figure 2.**
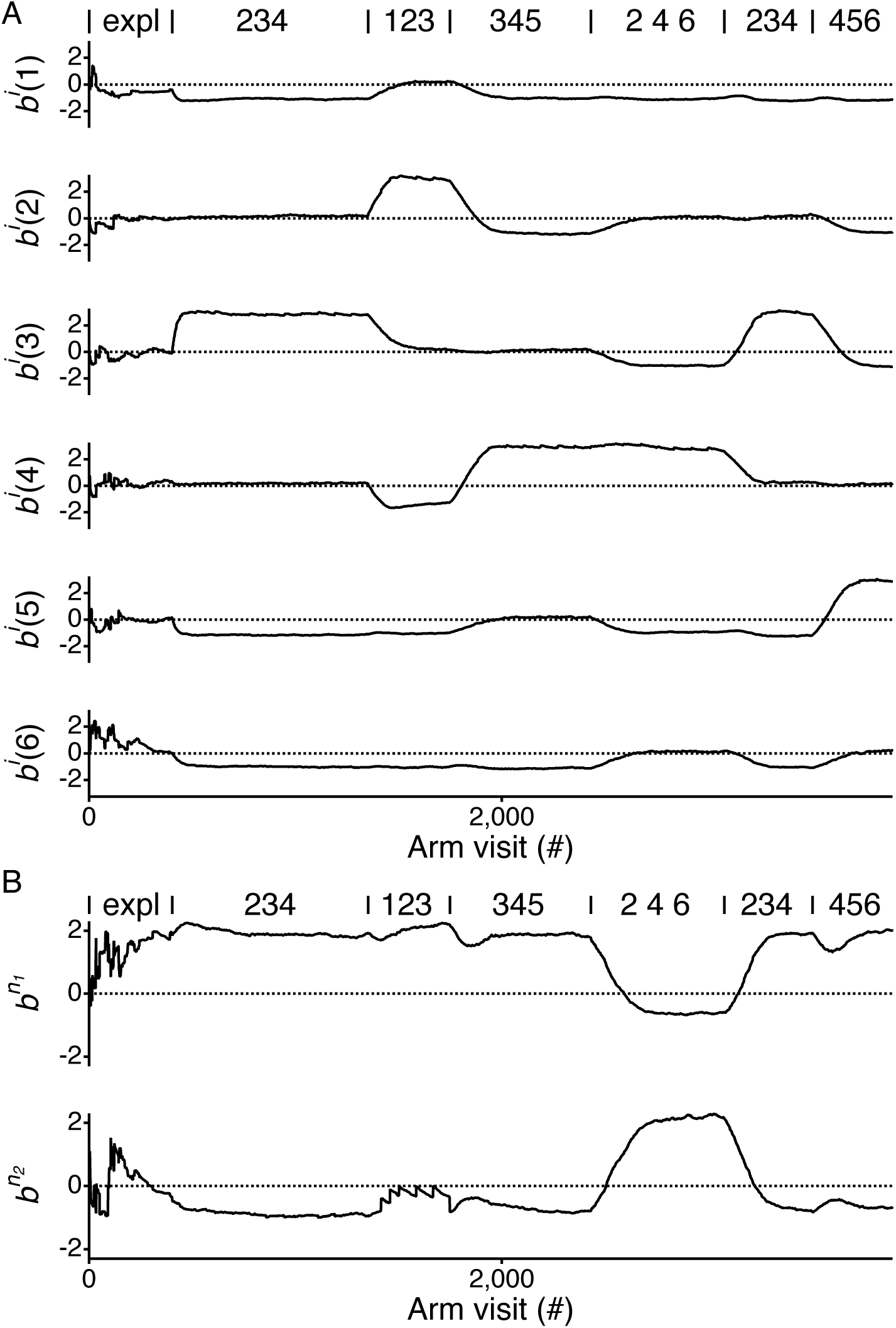
Dynamics of individual arm and neighbor preferences for the model. The average individual arm preferences (*β*^*α*^) (A) and neighbor arm preferences 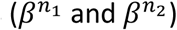 (B) across all contingencies and repeats of the model for the fit to the animal shown in Fig 5C. The values shown are those prior to passing through the exponential for the Softmax. Contingencies indicated as in Fig 1F.

**Supplementary Figure 3.**
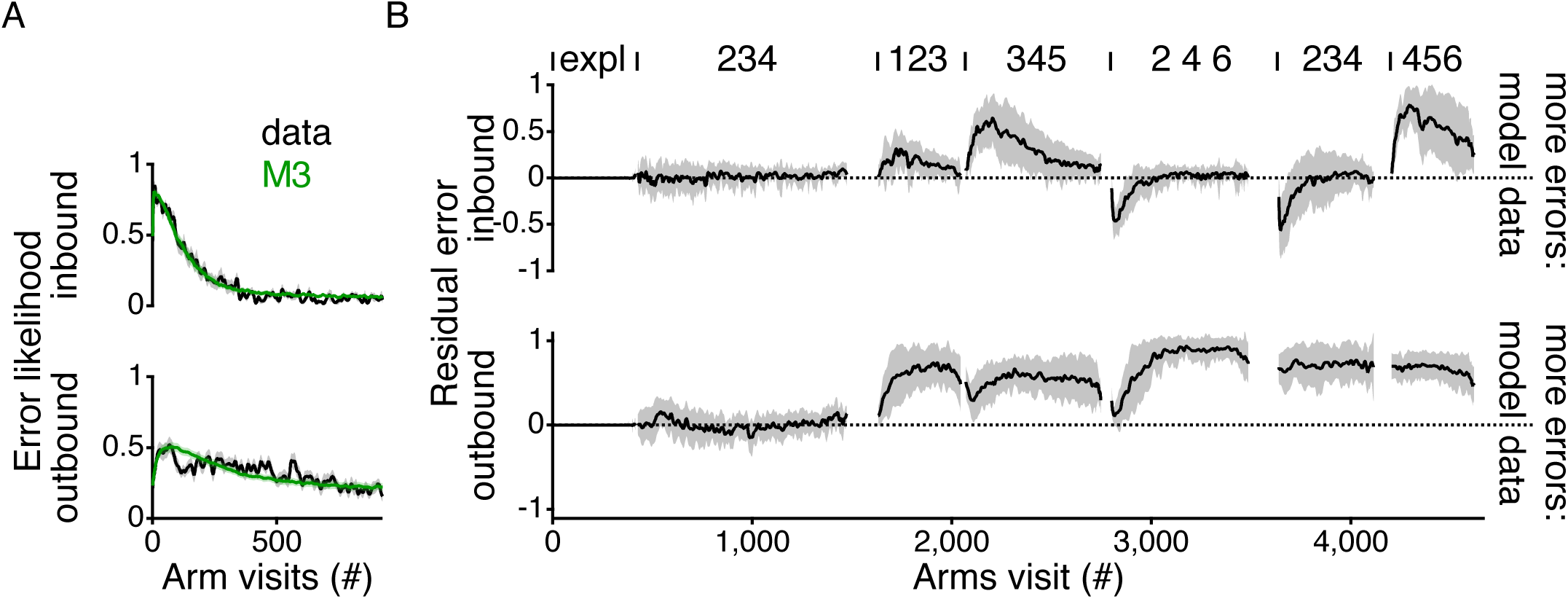
Fitting M3 to first contingency does not predict subsequent contingencies. (A) Average inbound and outbound errors for the data (black) and model M3 (green) after fitting M3 to each individual animal. (B) Average residuals between the fit to each individual animal and the model fit only to the first alternation contingency. See Fig. 6D for comparison.

**Supplementary Figure 4.**
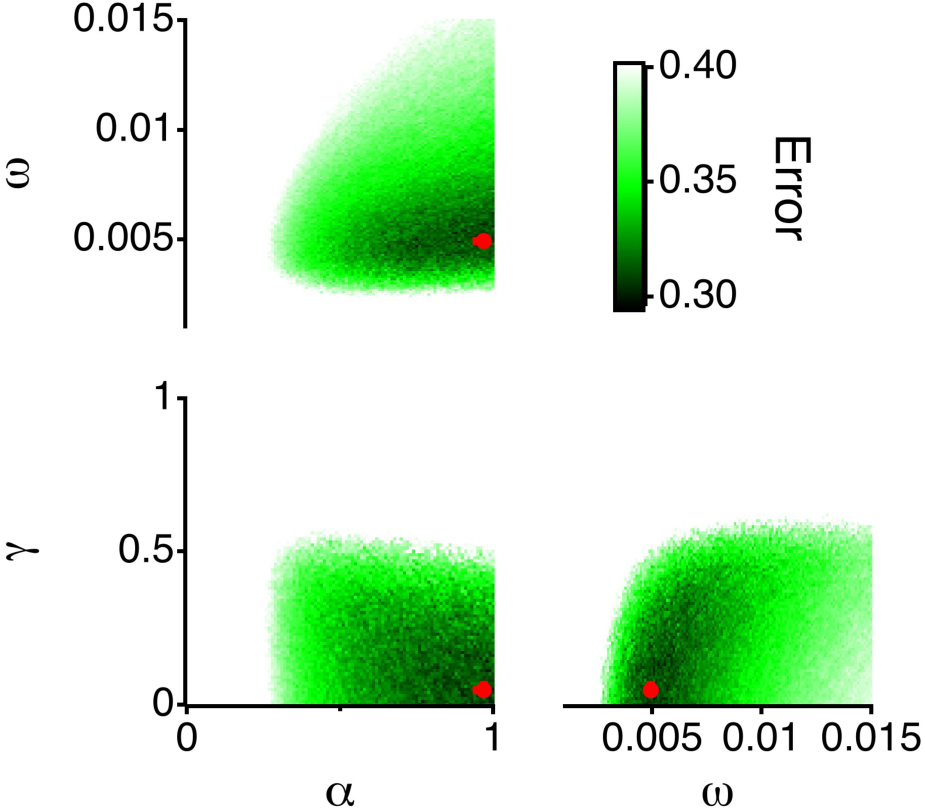
Error landscape for M3. (A) Three-dimensional space of parameters projected onto the plane for the parameters from the fit. For instance, for the *α*/*γ* plot, the plane for the fit value of *ω* is chosen. The median and interquartile range for the parameters for the same rat from Fig. 3 for 24 fits are plotted as the red dot with errors bars in both axes (obscured by the dot). The color scale in the background shows the error between the model and the data.

**Supplementary Figure 5.**
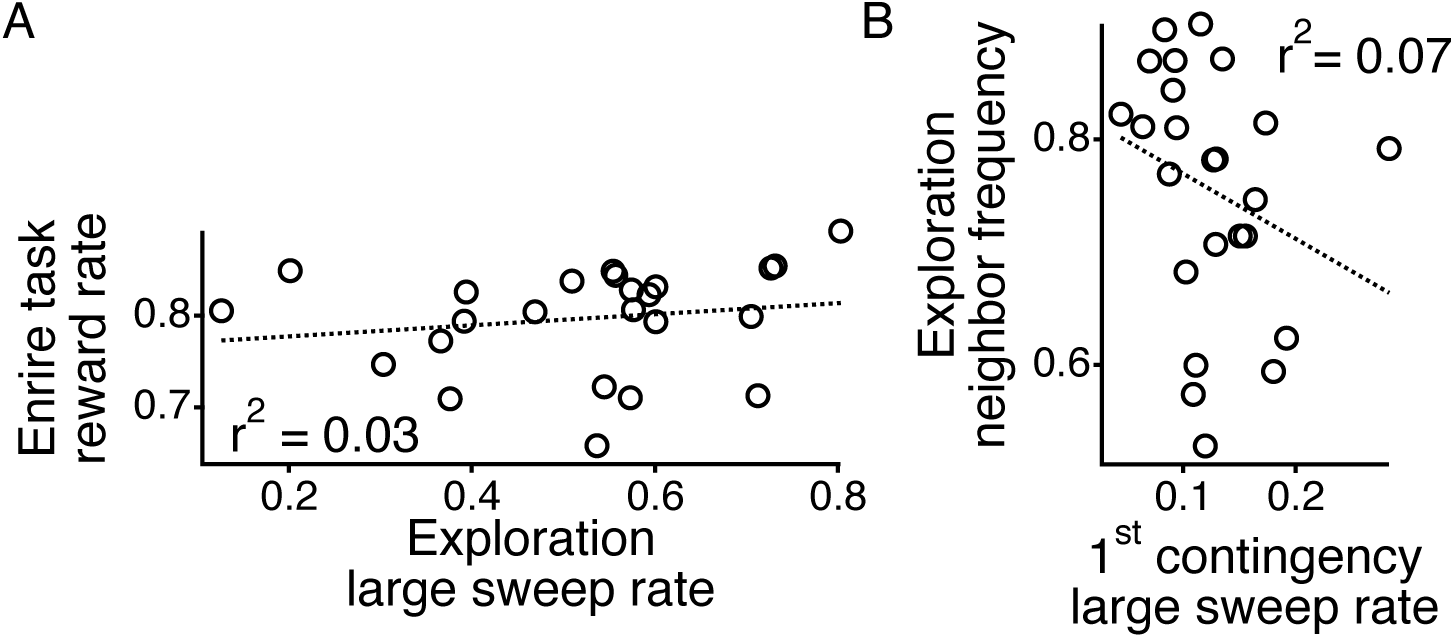
Lack of correlation between metrics. (A) Entire task reward rate plotted relative to the large sweep (>3 arms) rate during the exploratory period. Dotted line shows linear fit. (B) Neighbor transition frequency during the exploratory period plotted relative to the large sweep rate during the first alternation contingency for each animal. Dotted line shows linear fit.

**Supplementary Figure 6.**
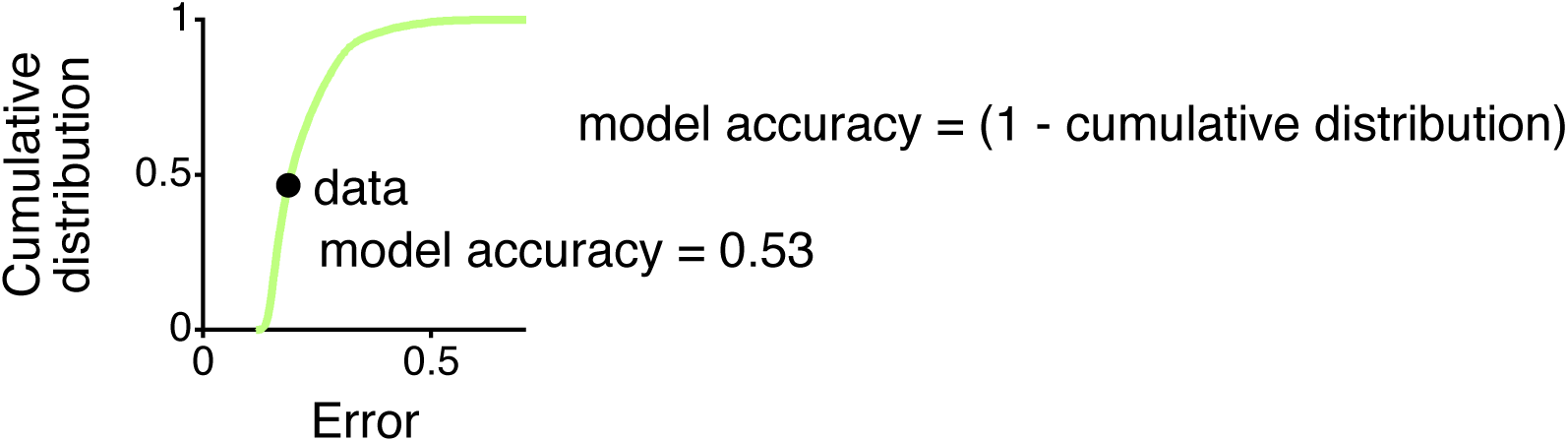
Calculation of model accuracy. Cumulative distribution for all of the RMS difference errors between individual simulations of the model and the 200 repeats of the model for the outbound errors of the fourth contingency for the animal displayed in Fig. 6A. The RMS difference error for the data is shown in the black circle. Model accuracy is the fraction of the cumulative distribution that falls to the right of the data.

## References

1. Sugrue, L. P., Corrado, G. S. & Newsome, W. T. Matching behavior and the representation of value in the parietal cortex. Science 304, 1782–1787 (2004).

2. Lau, B. & Glimcher, P. W. Dynamic response-by-response models of matching behavior in rhesus monkeys. J Exp Anal Behav 84, 555–579 (2005).

3. Samejima, K., Ueda, Y., Doya, K. & Kimura, M. Representation of action-specific reward values in the striatum. Science 310, 1337–1340 (2005).

4. Lee, M. D. & Wagenmakers, E. J. Bayesian Cognitive Modeling: A Practical Course. (Cambridge University Press, 2014).

5. Glascher, J. P. & O’Doherty, J. P. Model-based approaches to neuroimaging: combining reinforcement learning theory with fMRI data. Wiley Interdiscip Rev Cogn Sci 1, 501–510 (2010).

6. Ruediger, S. et al. Learning-related feedforward inhibitory connectivity growth required for memory precision. Nature 473, 514–518 (2011).

7. Ribeiro, M. et al. Meningeal *γδ* T cell-derived IL-17 controls synaptic plasticity and short­term memory. Sci Immunol 4, eaay5199 (2019).

8. Wang, F. et al. Myelin degeneration and diminished myelin renewal contribute to age-related deficits in memory. Nat Neurosci 23, 481–486 (2020).

9. Vasek, M. J. et al. A complement-microglial axis drives synapse loss during virus-induced memory impairment. Nature 534, 538–543 (2016).

10. Faraco, G. et al. Dietary salt promotes cognitive impairment through tau phosphorylation. Nature 574, 686–690 (2019).

11. Awasthi, A. et al. Synaptotagmin-3 drives AMPA receptor endocytosis, depression of synapse strength, and forgetting. Science 363, (2019).

12. Sigurdsson, T., Stark, K. L., Karayiorgou, M., Gogos, J. A. & Gordon, J. A. Impaired hippocampal-prefrontal synchrony in a genetic mouse model of schizophrenia. Nature 464, 763–767 (2010).

13. Mukai, J. et al. Recapitulation and Reversal of Schizophrenia-Related Phenotypes in Setd1a-Deficient Mice. Neuron 104, 471–487.e12 (2019).

14. Frank, L. M., Brown, E. N. & Wilson, M. A. Trajectory encoding in the hippocampus and entorhinal cortex. Neuron 27, 169–178 (2000).

15. Jadhav, S. P., Kemere, C., German, P. W. & Frank, L. M. Awake hippocampal sharp-wave ripples support spatial memory. Science 336, 1454–1458 (2012).

16. Shin, J. D., Tang, W. & Jadhav, S. P. Dynamics of Awake Hippocampal-Prefrontal Replay for Spatial Learning and Memory-Guided Decision Making. Neuron 104, 1110–1125.e7 (2019).

17. Fernandez-Ruiz, A. et al. Long-duration hippocampal sharp wave ripples improve memory. Science 364, 1082–1086 (2019).

18. Kastner, D. B., Gillespie, A. K., Dayan, P. & Frank, L. M. Memory alone does not account for the way rats learn a simple spatial alternation task. Journal of Neuroscience 40, 7311–7317 (2020).

19. Brunton, B. W., Botvinick, M. M. & Brody, C. D. Rats and humans can optimally accumulate evidence for decision-making. Science 340, 95–98 (2013).

20. Poddar, R., Kawai, R. & Olveczky, B. P. A fully automated high-throughput training system for rodents. PLoS One 8, e83171 (2013).

21. Rivalan, M., Munawar, H., Fuchs, A. & Winter, Y. An Automated, Experimenter-Free Method for the Standardised, Operant Cognitive Testing of Rats. PLoS One 12, e0169476 (2017).

22. International Brain Laboratory. An International Laboratory for Systems and Computational Neuroscience. Neuron 96, 1213–1218 (2017).

23. Sutton, R. S. & Barto, A. G. Reinforcement Learning. (MIT Press, 1998).

24. Silver, D. et al. Mastering the game of Go without human knowledge. Nature 550, 354–359 (2017).

25. Silver, D. et al. A general reinforcement learning algorithm that masters chess, shogi, and Go through self-play. Science 362, 1140–1144 (2018).

26. Jaderberg, M. et al. Human-level performance in 3D multiplayer games with population-based reinforcement learning. Science 364, 859–865 (2019).

27. Vinyals, O. et al. Grandmaster level in StarCraft II using multi-agent reinforcement learning. Nature 575, 350–354 (2019).

28. Gershman, S. J. & Niv, Y. Learning latent structure: carving nature at its joints. Curr Opin Neurobiol 20, 251–256 (2010).

29. Lake, B. M., Ullman, T. D., Tenenbaum, J. B. & Gershman, S. J. Building machines that learn and think like people. Behav Brain Sci 40, e253 (2017).

30. Sorge, R. E. et al. Olfactory exposure to males, including men, causes stress and related analgesia in rodents. Nat Methods 11, 629–632 (2014).

31. Singer, A. C. & Frank, L. M. Rewarded outcomes enhance reactivation of experience in the hippocampus. Neuron 64, 910–921 (2009).

32. Singer, A. C., Karlsson, M. P., Nathe, A. R., Carr, M. F. & Frank, L. M. Experience-dependent development of coordinated hippocampal spatial activity representing the similarity of related locations. Journal of Neuroscience 30, 11586–11604 (2010).

33. Kim, S. M. & Frank, L. M. Hippocampal lesions impair rapid learning of a continuous spatial alternation task. PLoS One 4, e5494 (2009).

34. Williams, R. J. Simple Statistical Gradient-Following Algorithms for Connectionist Reinforcement Learning. Machine Learning 8, 229–256 (1992).

35. Todd, M. T., Niv, Y. & Cohen, J. D. in Advances in Neural Information Processing Systems 21 (eds. Koller, D., Schuurmans, D., Bengio, Y. & Bottou, L.) 1689–1696 (Curran Associates, Inc., 2009).

36. Suri, R. E. & Schultz, W. A neural network model with dopamine-like reinforcement signal that learns a spatial delayed response task. Neuroscience 91, 871–890 (1999).

37. Li, J. & Daw, N. D. Signals in human striatum are appropriate for policy update rather than value prediction. Journal of Neuroscience 31, 5504–5511 (2011).

38. Zilli, E. A. & Hasselmo, M. E. Modeling the role of working memory and episodic memory in behavioral tasks. Hippocampus 18, 193–209 (2008).

39. Lloyd, K., Becker, N., Jones, M. W. & Bogacz, R. Learning to use working memory: a reinforcement learning gating model of rule acquisition in rats. Front Comput Neurosc 6, 87 (2012).

40. Lintusaari, J., Gutmann, M. U., Dutta, R., Kaski, S. & Corander, J. Fundamentals and Recent Developments in Approximate Bayesian Computation. Syst. Biol. 66, e66–e82 (2017).

41. Luksys, G., Gerstner, W. & Sandi, C. Stress, genotype and norepinephrine in the prediction of mouse behavior using reinforcement learning. Nat Neurosci 12, 1180–1186 (2009).

42. Daw, N. D. in Decision Making, Affect, and Learning: Attention and Performance XXIII 3–38 (Oxford University Press, 2011). doi:10.1093/acprof:oso/9780199600434.003.0001

43. Akam, T., Costa, R. & Dayan, P. Simple Plans or Sophisticated Habits? State, Transition and Learning Interactions in the Two-Step Task. PLoS Comput. Biol. 11, (2015).

44. Kirkpatrick, J. et al. Overcoming catastrophic forgetting in neural networks. Proc Natl Acad Sci USA 114, 3521–3526 (2017).

45. Zenke, F., Ben Poole & Ganguli, S. Continual learning through synaptic intelligence. 3987–3995 (2017).

46. Galsworthy, M. J., Paya-Cano, J. L., Monleon, S. & Plomin, R. Evidence for general cognitive ability (g) in heterogeneous stock mice and an analysis of potential confounds. Genes Brain Behav. 1, 88–95 (2002).

47. Matzel, L. D. et al. Individual differences in the expression of a ‘general’ learning ability in mice. J Neurosci 23, 6423–6433 (2003).

48. Matzel, L. D. et al. Exploration in outbred mice covaries with general learning abilities irrespective of stress reactivity, emotionality, and physical attributes. Neurobiol Learn Mem 86, 228–240 (2006).

